# Synthesis of mRNA lipid nanoparticles for engineering GD2 CAR T and CAR NK cells against neuroblastoma

**DOI:** 10.64898/2025.12.15.692195

**Authors:** Chih-Chun Chang, Lei Shi, Soon H. Choi, Andrea Pennati, Vasiliki Valkanioti, Christian M. Capitini, Sandro Mecozzi, Jacques Galipeau

**Author notes:** Correspondence should be addressed to J.G. and S.M. ( &).

## Abstract

Engineering T and natural killer (NK) cells with chimeric antigen receptors (CAR) creates effective adoptive cell transfer therapies for cancer treatment. However, using viral transduction as a primary genetic modification method adds regulatory burdens and is expensive to produce at scale. Delivering mRNA encoding CAR via lipid nanoparticles (LNPs) has been explored as a potent non-viral method to generate CAR immune cells. Still, it has not been optimized for CAR treatment of neuroblastoma to date. An LNP formulation to deliver mRNA encoding a GD2 CAR into human T and NK cells was designed and characterized by dynamic light scattering for size distribution, z-average diameter, polydispersity index, and ζ potential. Fluorescent reporter detection persisted for more than 1 week after mRNA LNP transfection, without affecting T or NK cell viability. The potency of GD2 CAR T cells with 79.9% reporter positivity and GD2 CAR NK cells with 26.6% reporter positivity was assessed *in vitro* against the GD2^+^ neuroblastoma cell line CHLA20. GD2 CAR T or CAR NK cells could effectively target and kill neuroblastoma cells in a dose-dependent fashion, and GD2 CAR T cells showed increased IFNγ production. This study shows mRNA LNPs are a promising non-viral approach for generating GD2 CAR T and CAR NK cells, potentially offering a safer and more cost-effective alternative to current viral vector-based methods.

## Introduction

One promising strategy for generating chimeric antigen receptor (CAR) T cells and CAR NK cells is the transfection of cells via the delivery of CAR-encoded mRNA.^1–5^ This non-viral approach allows for transient expression of the CAR, reduces regulatory concerns around monitoring for replication competence and insertional oncogenesis, and somewhat mitigates the financial expense associated with viral vector-based methods.^6–8^ The use of mRNA for CAR cell therapies has demonstrated therapeutic efficacy against various solid tumors and hematologic malignancies in both preclinical studies and clinical trials.^1,2^ Electroporation facilitates mRNA entry into cells by creating temporary disruptions in the cell membrane; however, this technique can induce membrane damage, transient decreases in cell viability, loss of cellular content, and changes in gene and protein expression.^9–11^ These challenges highlight the need for improved delivery methods for mRNA-based CAR cell therapies.

Lipid nanoparticles (LNPs) are promising alternatives to electroporation for mRNA delivery in CAR cell engineering.^10,12,13^ Previous studies have demonstrated effective transfection in various immune cell types, including myeloid and lymphoid cells, with reduced cytotoxicity compared to electroporation.^12–16^ LNPs are more clinically advanced for RNA delivery than other types of delivery systems, as evidenced by FDA approvals for RNA-based therapeutics such as patisiran and COVID-19 vaccines.^17–19^ Typically, mRNA LNPs consist of four main components: an ionizable lipid, cholesterol, phospholipid, and lipid-anchored polyethylene glycol (PEG-lipid).^20,21^ The ionizable lipid complexes with negatively charged mRNA via electrostatic interaction enable the encapsulation of nucleic acids.^22^ One proposed mechanism for mRNA release from endosomes involves an interaction between ionizable lipids in the LNP and endosomal lipids, resulting in membrane destabilization and rupture.^23^ Cholesterol and phospholipids provide structural support, stabilizing the LNP under physiological conditions and facilitating membrane fusion during endosomal escape.^24,25^ The PEG-lipid coating provides a hydrophilic layer that confers stealth properties, preventing recognition by the mononuclear phagocyte system and particle aggregation.^25,26^

Recent studies have shown that LNP-mediated delivery of mRNA encoding CD123- and CD19-specific CARs significantly outperforms electroporation methods in CAR-T performance against leukemia cell lines.^10,13^ Additionally, mRNA LNP technology excels at generating functional BCMA and CD19 CAR NK cells with high antitumor activity.^3^ Building on these findings, we developed an LNP formulation using Food and Drug Administration (FDA)-approved lipids to deliver mRNA encoding a second-generation CAR targeting GD2, which is overexpressed in cancers like neuroblastoma^27^ and is a promising target for CAR T therapy in early-phase trials.^28–33^ We successfully generated GD2 CAR T and CAR NK cells through transfection with an mRNA LNP, and validated their potency against GD2^+^ neuroblastoma cells.

## Results

We developed LNPs using FDA-approved lipid components for delivering mRNA to human T and NK cells. The formulation consisted of an ionizable lipid, a helper lipid, cholesterol, and a PEGylated lipid. To formulate the mRNA LNP, all lipid components were dissolved in ethanol as the organic phase, and mRNA was prepared in citrate buffer as the aqueous phase. The mRNA LNP was spontaneously formed by rapidly mixing the two phases with a pipette. After formation, the LNPs were purified and concentrated through dialysis and centrifugal filtration (**Figure 1A**).^34^ We compared blank LNPs with mRNA-containing LNPs prepared using two different mRNA cargos synthesized via *in vitro* transcription (IVT) from DNA templates (**Figure S1**). The LNPs were loaded with either eGFP-T2A-mApple (GTA) mRNA or GD2 CAR-T2A-mCherry (CTC) mRNA. Characterization of the concentrated LNPs included size distribution, z-average diameter, polydispersity index (PDI), and ζ potential using dynamic light scattering (DLS). All LNPs displayed a monodisperse profile, with z-average sizes under 100 nm and a PDI below 0.2, indicating a homogeneous particle population (**Figure 1B,C**).^35^ Despite CTC mRNA being approximately 750 bases longer than GTA mRNA, the mRNA cargo sizes did not significantly influence LNP sizes, consistent with previous findings.^36,37^ The surface charges of LNPs were slightly negative, ranging from -5 to -10 mV, a value considered approximately neutral (**Figure 1D**).^38–40^ Relative encapsulation efficiencies (EE%) of GTA and CTC mRNA were over 95%, determined by a RiboGreen assay (**Figure S2**). The apparent p*K*a of our LNP was 6.83, well within the optimal range for potent mRNA LNPs that can be effectively ionized during endosomal maturation (**Figure 1E**).^41^ To assess cell viability following LNP transfection, Jurkat cells were treated with blank LNPs at escalating doses of total lipids or with mRNA LNPs at the same lipid doses, each containing the corresponding amount of mRNA (**Figure 1F,G**). Both blank and mRNA LNPs had minimal impact on Jurkat cell viability, indicating the ideal biocompatibility of the lipid choices.

**Figure 1.**
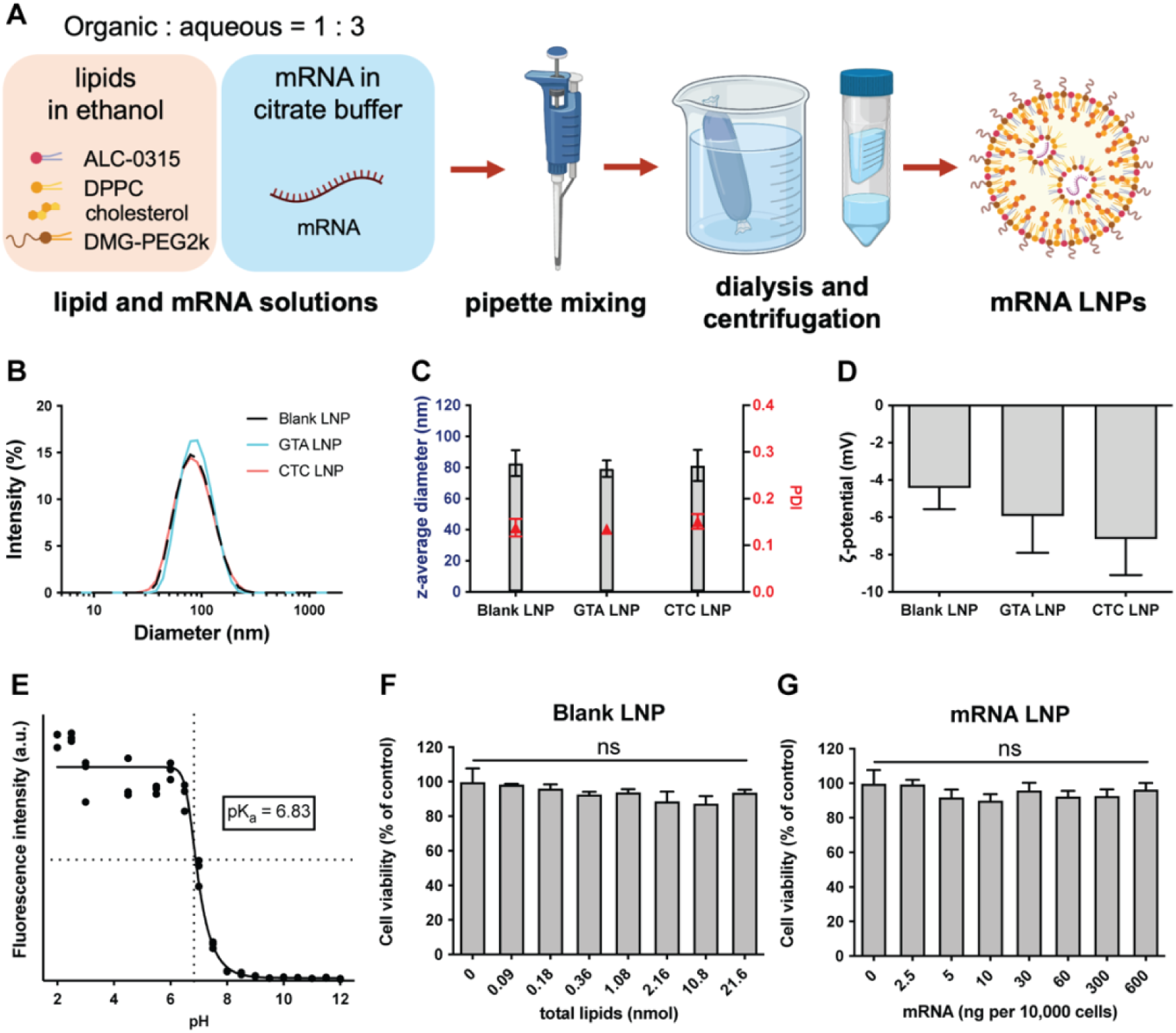
Preparation and characterization of blank and mRNA LNPs (A) Key lipid components include ionizable lipids (ALC-0315), helper lipids (DPPC), cholesterol, and PEGylated lipids (DMG-PEG2000), which were dissolved in ethanol as an organic phase. Citrate buffer was used to prepare the mRNA aqueous solution. Self-assembly of LNP was facilitated by pipette mixing of the two phases, followed by dialysis and centrifugal filtration. The scheme was created with BioRender.com. (B) Size distribution (C) Z-average diameter, polydispersity index (PDI), and (D) ζ potential of the LNP loaded with eGFP-T2A-mApple (GTA) or CAR-T2A-mCherry mRNA (CTC) and the LNP without mRNA (Blank) were determined by DLS. Data are presented as mean ± standard deviation (*n* = 3). (E) TNS assay plot of mRNA LNPs demonstrating the ionizability of the particle surface. (F) The cell viability of Jurkat treated with the blank LNP at 0.09, 0.18, 0.36, 1.08, 2.16, 10.8, and 21.6 nmol of total lipids per 10,000 cells for 24 h showed minimal cytotoxicity associated with lipid components. (G) The cell viability of Jurkat cells was assessed after treatment with mRNA-loaded LNP containing the corresponding amount of mRNA per 10,000 cells, with total lipids matching the doses of the blank LNP. Viability was normalized to untreated cells. Data are presented as mean ± standard deviation (*n* = 3). ns, not significant, unpaired two-sided Mann-Whitney test.

To evaluate the efficiency of LNP-based mRNA delivery, enhanced green fluorescent protein (eGFP) and mApple were selected as reporter proteins encoded by eGFP-T2A-mApple mRNA. The green and red fluorescence signals correlate with the transfection efficiency of the mRNA LNP, as a measure of functional mRNA delivery. The delivery of eGFP-T2A-mApple mRNA was first assessed in the immortalized cell lines Jurkat and NK-92. These cells were treated with the GTA LNP at 300 ng per 10,000 cells (**Figure 2A,B; Figure S3**). Both cell lines demonstrated approximately 90% reporter-positive cells, without affecting cell viability. The overlap of eGFP and mApple fluorescence in the images confirmed the successful function of the T2A peptide, resulting in simultaneous expression of both fluorescent reporters, and the expression of eGFP and mApple in Jurkat cells persisted for over 10 days (**Figure 2C**).

**Figure 2.**
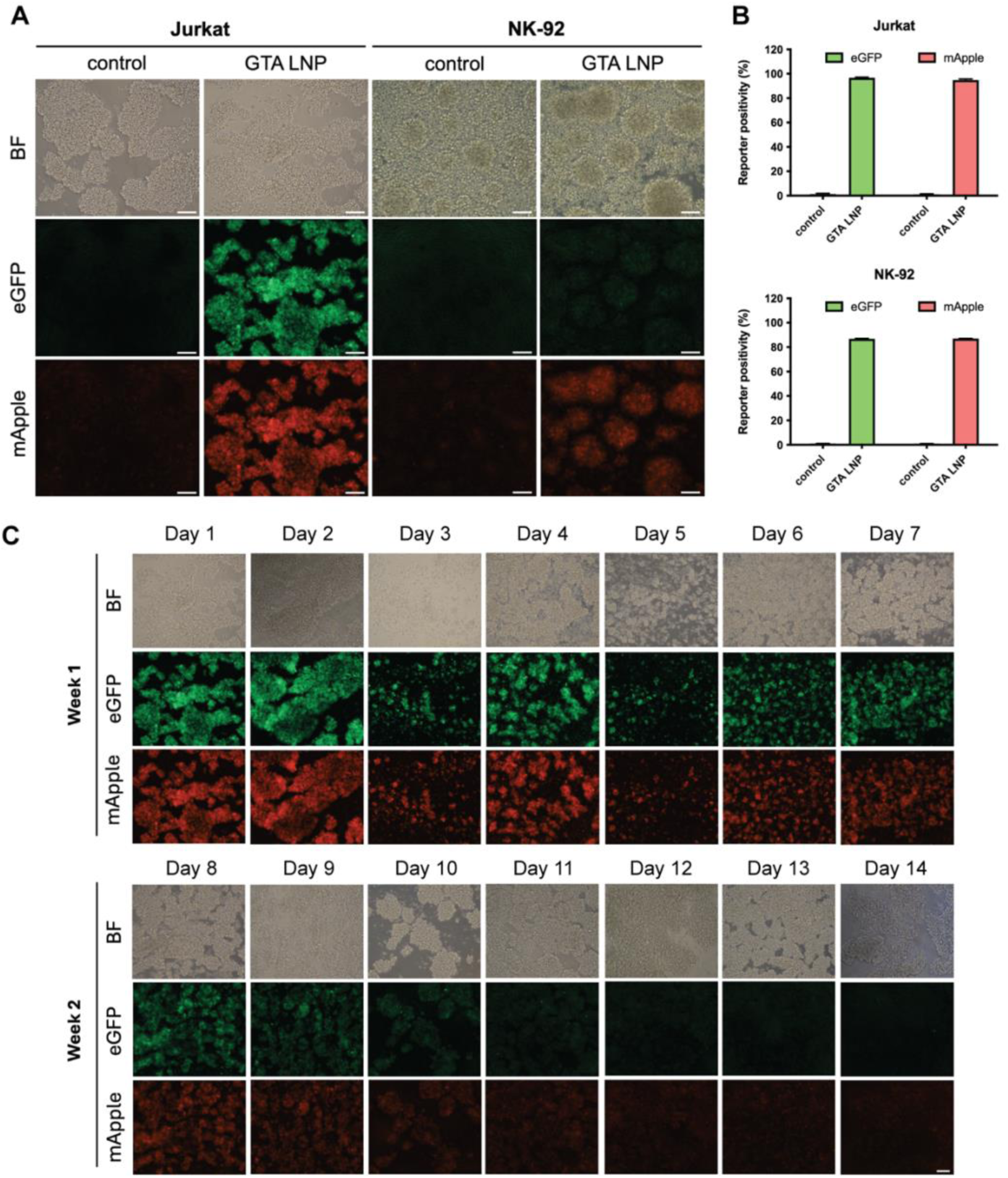
GTA mRNA delivery into Jurkat and NK-92 cells by LNPs Jurkat and NK-92 cells were transfected for 24 h with the GTA LNP at 300 ng mRNA per 10,000 cells. (A) Representative bright-field (BF) and fluorescent images (eGFP and mApple) of the Jurkat and NK-92 cells. Scale bar = 100 μm. (B) Reporter positivity of the LNP-transfected Jurkat and NK-92 cells was analyzed via flow cytometry. Data are presented as mean ± standard deviation (*n* = 3). (C) Representative images of GTA LNP-transfected Jurkat cells showing eGFP and mApple expression trend for 2 weeks. Scale bar = 100 μm.

After observing efficient transfection in Jurkat and NK-92 cells, a transfection study was conducted using murine T cells isolated from splenocytes and cultured in R10 medium supplemented with recombinant human interleukin-2 (rhIL-2) (**Figure S4A-D**). Murine T cells were activated using Dynabeads Mouse T-activator at a 1:1 bead-to-cell ratio, as CD3/CD28 activation significantly enhances LNP-mediated transfection.^42^ After one day of activation, the cells were transfected with the GTA LNP at 30 ng or 60 ng mRNA per 10,000 cells. Murine T cells were cultured at 1 million cells/mL, ten times the density of the Jurkat culture. The high cell density is necessary for effective primary T-cell activation and transfection.^43^ To prevent overdilution of the cell culture medium with LNP solutions, the amount of mRNA per 10,000 cells was reduced to 30 or 60 ng for the subsequent primary cell studies. Our findings suggest that the GTA LNP is not species-specific and can also transfect metabolically active murine T cells.

We next sought to confirm whether the GTA LNP can transfect primary human T cells. We isolated primary T cells from healthy donor peripheral blood mononuclear cells (PBMCs) and activated them for 72 h using a CD3 antibody-coated plate and medium supplemented with CD28 antibodies and rhIL-2. Results from fluorescent imaging and flow cytometry confirmed that the GTA LNP effectively induced the expression of the reporter proteins in human T cells, while free mRNA or blank LNP groups did not show the expression (**Figure 3A-D; Figure S5**). Viability analysis indicated minimal toxicity from the GTA LNP, with over 90% of primary human T cells remaining alive after transfection (**Figure 3B**). The percentage of cells expressing the mApple reporter was 44.1% and 52.5% for the 30 ng and 60 ng mRNA doses, respectively. In contrast, the proportion of cells expressing the eGFP reporter was higher, at 87.1% and 88.1% for the low and high doses, respectively (**Figure 3D**). To analyze the kinetics of reporter protein expression, *ex vivo*-modified primary human T cells were monitored daily for 8 days using both fluorescence imaging and flow cytometry (**Figure 3E,F; Figure S6**). The expression of eGFP and mApple reached their maximum levels at 24 hours post-transfection. Notably, eGFP positivity in cells transfected with 60 ng mRNA per 10,000 cells remained around 80% throughout 8 days, while eGFP positivity in the 30 ng mRNA dose group gradually decreased from 83% to 57% throughout the same period. In contrast, the mApple positivity exhibited an exponential decay immediately after its expression peak for both doses. The discrepancy between eGFP and mApple expression underscores the different kinetics of the two reporters.

**Figure 3.**
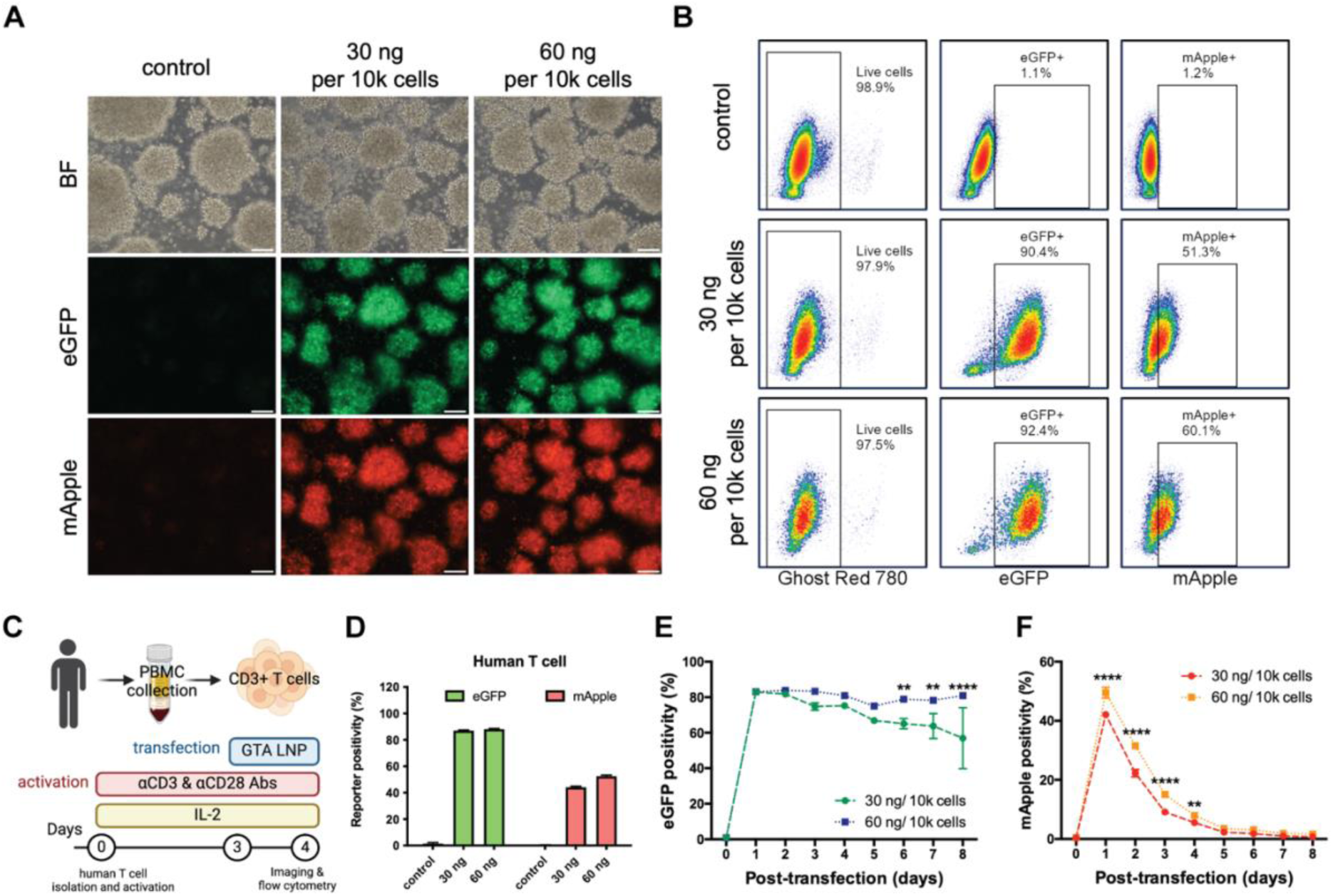
GTA mRNA delivery into primary human T cells by LNPs (A) Representative bright-field (BF) and fluorescent images of the human T cells with or without LNP transfection. Images were taken 24 h after transfection. Upon activation, human T cells tend to form cellular clusters. Scale bar = 100 μm. (B) Representative flow cytometric density plot of Ghost Dye Red staining and fluorescent reporter expression of the human T cells under various treatments. (C) Experimental design for *ex vivo* human T cell transfection. Primary human T cells were isolated from PBMCs donated by healthy volunteers and supplemented with 150 IU/mL rhIL-2 every other day. The isolated T cells were immediately activated for 72 hours with αCD3 and αCD28 antibodies before LNP transfection. (D) Reporter positivity of the non-transfected control and the LNP-transfected human T cells. Data are presented as mean ± standard deviation (*n* = 3). (E)(F) eGFP and mApple expressed by the transfected cells were monitored for 8 days using flow cytometry. Data are presented as mean ± standard deviation (*n* = 3). **p < 0.01; ****p < 0.0001, two-way ANOVA with Sidak’s multiple comparison test. The scheme was created with BioRender.com.

We next investigated the role of apolipoprotein E4 (ApoE4) in the cellular uptake of the CTC LNP, given its known importance in facilitating endocytosis through the low-density lipoprotein receptor pathway. The mRNA delivery using CTC LNP to primary T cells was tested with or without additional human ApoE4 (1 μg/mL) to determine if ApoE4 was necessary for human T-cell transfection under serum-containing conditions (**Figure S7**).^10^ The mCherry positivity was approximately 80% for both doses, regardless of the presence or absence of human ApoE4 when using serum-containing medium for transfection. Various bovine-origin apolipoprotein isoforms in the serum-containing medium may also facilitate PEG shedding and the formation of a protein corona around our LNPs^44^ thereby inducing structural and compositional changes in LNPs and promoting the release of mRNA cargo into the cytoplasm.^45^ Therefore, we decided to proceed using serum-containing medium without human ApoE4 for subsequent CTC LNP transfection experiments.

To generate GD2 CAR T cells, 14G2a-41BB-CD3ζ (GD2 CAR)-T2A-mCherry mRNA was encapsulated into the LNP and used to transfect activated T cells isolated from three healthy donors (**Figure 4A**). The varying transfection efficiencies revealed donor-to-donor variability, with one donor showing lower reporter expression. Although mCherry positivity varied among cells from different donors, cell viability remained above 90% after treatment with the CTC LNP (**Figure 4B**). To directly determine GD2 CAR expression levels, a c-Myc tag was inserted between the single-chain variable fragment (scFv) and the hinge region of the CAR construct. CAR surface expression was detected by anti-c-Myc in 69.8% and 45.7% of cells in the low and high LNP dose groups, respectively, while mCherry reporter expression was detected in 79.9% and 69.9% of cells (**Figure 4C-E**). These results indicated that the correlation between mRNA LNP potency and increasing doses was not always linear and might plateau at higher concentrations.^46^ Remarkably, we successfully engineered primary human T cells with GD2 CAR using the CTC LNP, as evidenced by the mCherry signal and c-Myc positivity. Swapping LNP cargo from a reporter mRNA to CAR mRNA had minimal impact on T cell viability.

**Figure 4.**
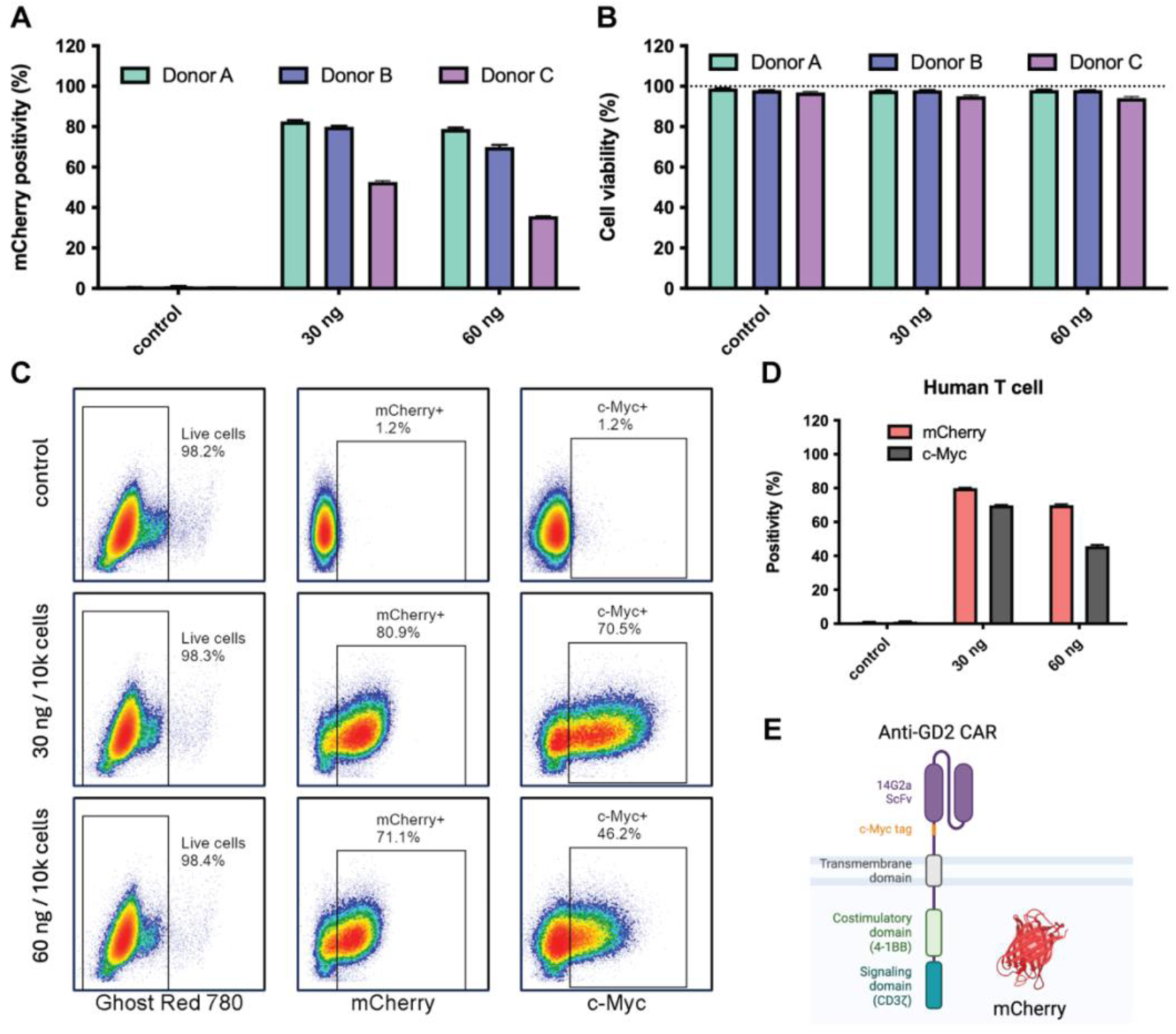
Anti-GD2 CAR and mCherry expression of primary human T cells Primary human T cells were treated with the CTC LNP containing the corresponding doses of the mRNA for 24 h. (A)(B) mCherry positivity and cell viability for each donor and dose. Data are presented as mean ± standard deviation (*n* = 3). (C) Representative flow cytometric density plot of viability staining, mCherry, and c-Myc expression. (D) mCherry reporter and c-Myc positivity of the transfected T cells and non-transfected control. Data are presented as mean ± standard deviation (*n* = 3). (E) Schematic diagram of the GD2 CAR construct. After translation, mCherry remained within the cytosol, while the GD2 CAR was translocated to the T cell surface.

To evaluate LNP-mediated mRNA delivery on primary NK cells, we first isolated CD56^+^ NK cells from PBMCs using negative selection. The isolated NK cells were then expanded with irradiated K562 feeder cells expressing membrane-bound IL-15 and 4-1BB ligand (K562-mbIL15-41BBL) and transfected with the CTC LNP (**Figure 5A**). We analyzed the cell cycle progression of the expanded NK cells from separate donors on Days 3, 6, and 8 using propidium iodide (PI) staining (**Figure S8**). Flow cytometric analysis revealed distinct cell cycle distribution patterns among NK cells from three donors during *ex vivo* expansion. All donors’ NK cells exhibited a prominent G0/G1 peak with elevated proportions of cells in S phase on Day 3, indicating active DNA synthesis upon stimulation by feeder cells. Cell cycle distribution demonstrated donor-dependent variations correlated with cell expansion outcomes (**Figure 5B**). NK cells derived from Donors A and B showed a progressive decline in S-phase proportions on Day 6 and Day 8, alongside an increase in the G0/G1 fraction, correlating with increased cell numbers. In contrast, NK cells from Donor C exhibited weak proliferation in response to feeder cell activation. The impaired expansion capacity was indicated by persistently high proportions in S phase over time and a lack of significant increases in G0/G1 populations. The result suggested a potential G2/M arrest, where cells complete DNA synthesis but fail to progress through mitosis. To evaluate if NK expansion capacity influences the efficiency of genetic modification using mRNA LNP, we transfected the expanded NK cells after 8 days of incubation with feeder cells (**Figure 5C; Figure S9A**). Transfection efficiency was assessed 24 hours post-transfection with the GTA LNP at 30 ng mRNA per 10,000 cells using flow cytometry. NK cells from Donors A and B, which demonstrated robust expansion (cell numbers increased by more than 4-fold by Day 8), showed superior transfection efficiencies (Donor A: 89.4% eGFP^+^, 86.6% mApple^+^; Donor B: 84.5% eGFP^+^, 79.4% mApple^+^). Conversely, the poorly expanded culture from Donor C (cell numbers increased approximately 1.5-fold by Day 8) contained only about 10% eGFP^+^ or mApple^+^ cells. These results indicated a correlation between NK cell expansion and subsequent mRNA LNP transfection efficiency. To characterize the expression duration of the fluorescent reporter proteins, GTA LNP-transfected NK cells from Donor A were cultured for one week and harvested at designated time points for flow cytometric analysis (**Figure S9B**). The fluorescent protein positivity exceeded 85% on day 1 post-transfection for both eGFP and mApple reporters, and expression persisted for more than 1 week. Having established the relationship of NK cell expansion and transfection, we investigated the impact of LNP formulation (with or without GTA mRNA cargo) on NK cell phenotypes to evaluate its potential as a delivery system for NK cell-based immunotherapies (**Figure 5D-G**). The effects of LNPs on NK cell viability and phenotype were assessed using flow cytometry. Live cell gating demonstrated comparable viability across all conditions, suggesting high compatibility of both blank and mRNA-loaded LNPs with human NK cells (**Figure 5E**). CD3 and CD56 expression confirmed a similar proportion of CD3^−^CD56^+^ NK population among live cells with or without transfection. However, the expression of NKp46, an NK cell cytotoxicity receptor, decreased following LNP treatment (**Figure 5F**). Specifically,

**Figure 5.**
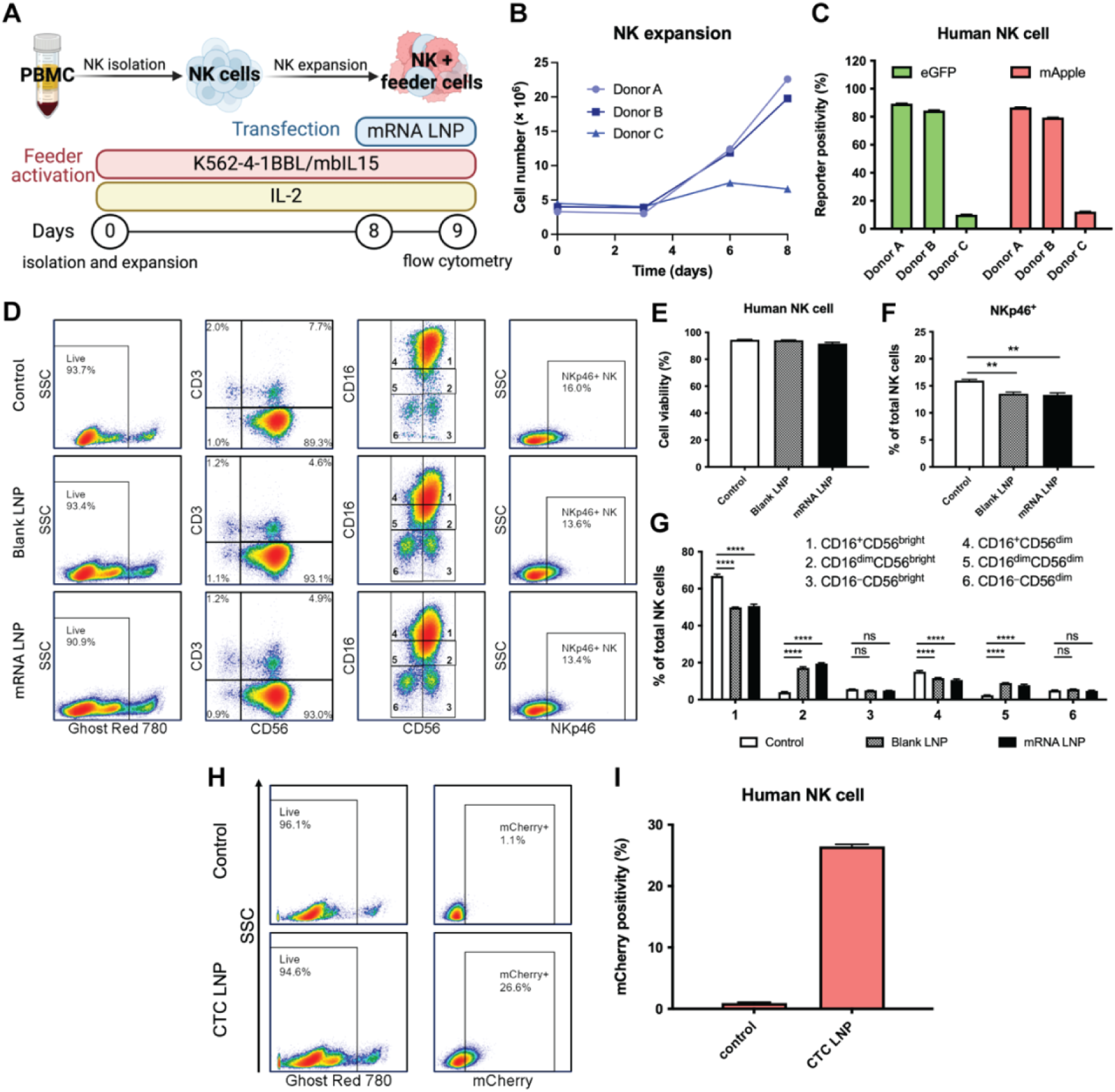
NK cell expansion and mRNA LNP transfection (A) Experimental design for *ex vivo* human NK cell expansion and transfection. Primary human NK cells were isolated from PBMCs donated by healthy volunteers and supplemented with 100 IU/mL rhIL-2 every other day. The isolated NK cells were immediately expanded and activated with irradiated K562-mbIL15-41BBL feeder cells before LNP transfection on Day 8. (B) NK expansion results of 3 separate donors. The cell numbers were counted with Trypan Blue before (Day 0) and after the feeder cell activation (Days 3, 6, and 8). (C) Reporter positivity of the LNP-transfected NK cells obtained from separate donors. The cells were transfected with GTA LNP (30 ng mRNA per 10,000 cells). (D) Representative flow cytometry gating for expanded NK cells treated with blank LNP, mRNA LNP (GTA mRNA), or without LNP treatment. The blank LNP matched the mRNA LNP in terms of total lipids added to the NK cells (Total lipids: 1.08 nmol per 10,000 cells, mRNA: 30 ng per 10,000 cells). The Ghost Red 780 was used for live cell gating. The lower right quadrant of the CD3 and CD56 plot represented the NK cell population (CD3^−^CD56^+^), which was gated for NK cell subsets and NKp46 expression. (E) Human NK cell viability was quantified based on Ghost Red staining for each treatment group by flow cytometry. (F) NKp46-positive cells were quantified for each treatment group by flow cytometry. (G) Distribution of NK cell subsets defined by CD16 and CD56 expression: 1. CD16^+^CD56^bright^, 2. CD16^dim^CD56^bright^, 3. CD16^−^CD56^bright^, 4. CD16^+^CD56^dim^, 5. CD16^dim^CD56^dim^, and 6. CD16^−^CD56^dim^ NK cells. (H) Representative flow cytometric density plot of viability staining and mCherry expression of non-transfected control and CTC LNP-treated group. (I) mCherry reporter positivity of the transfected NK cells and the non-transfected control. Data are presented as mean ± standard deviation (*n* = 3). ns, not significant; **p < 0.01; ****p < 0.0001, two-way ANOVA with Dunnett’s multiple comparison test. The scheme was created with BioRender.com.

NKp46^+^ NK cells decreased from 16% in the control to 13.6% in the blank LNP and 13.4% in the mRNA LNP. The subpopulations of NK cells based on CD16 and CD56 expression levels were further analyzed due to their distinct functional properties (**Figure 5D,G**).^47^ Our findings indicated a decrease in cytokine-producing CD16^+^CD56^bright^ NK cells from 66.8% in the control to 49.8% in the blank LNP group and 50.6% in the mRNA LNP group. On the other hand, an increase in CD16^dim^Cd56^bright^ NK cells from 4.0% in the control to 17.2% in the blank LNP group and 19.5% in the mRNA LNP group. Further investigation will be required to understand the mechanisms of interplay between LNP lipid components and NK phenotype shift. To generate GD2 CAR NK cells, Donor A’s NK cells were transfected with the CTC LNP at 30 ng mRNA per 10,000 cells following the same expansion and transfection procedure (**Figure 5H,I**). Flow cytometry showed 26.6% mCherry positivity within live NK cells, indicating successful CAR mRNA delivery to NK cells.

After generating GD2 CAR T and CAR NK cells using the CTC LNP, we investigated their potency against the GD2^+^ CHLA20 neuroblastoma cell line *in vitro*. CHLA20 cells expressing GFP were co-incubated with either non-transfected primary immune effector cells or CTC mRNA-transfected immune effector cells at different effector-to-target (E:T) ratios. Over approximately four days, the growth of CHLA20 cancer cells in both the CHLA20-only and co-culture groups was monitored at 4-hour intervals using the IncuCyte live imaging system (**Figure 6A; Figure S10**). Although GD2 CAR^+^ cell numbers were not normalized by reporter positivity (mCherry^+^ T cells: 79.9%; mCherry^+^ NK cells: 26.6%) in the IncuCyte assay, the GD2 CAR groups showed significantly better anti-tumor activity against GFP^+^ CHLA20 neuroblastoma cells than compared to non-transfected primary cell counterparts at effector-to-target (E:T) ratios of 4:1 and 2:1, but no differences in cytotoxicity at the 1:1 ratio (**Figure 6B**). To examine GD2 CAR T cell activation, supernatants from the IncuCyte plate were collected and analyzed using an IFN-γ ELISA assay (**Figure 6C**). The CAR T cells produced significantly more IFN-γ than unmodified primary T cells. This enhanced cytokine secretion, in combination with the observed cytotoxicity against GD2^+^ neuroblastoma, suggests that transfecting T cells with mRNA LNPs can generate potent CAR T cells.

**Figure 6.**
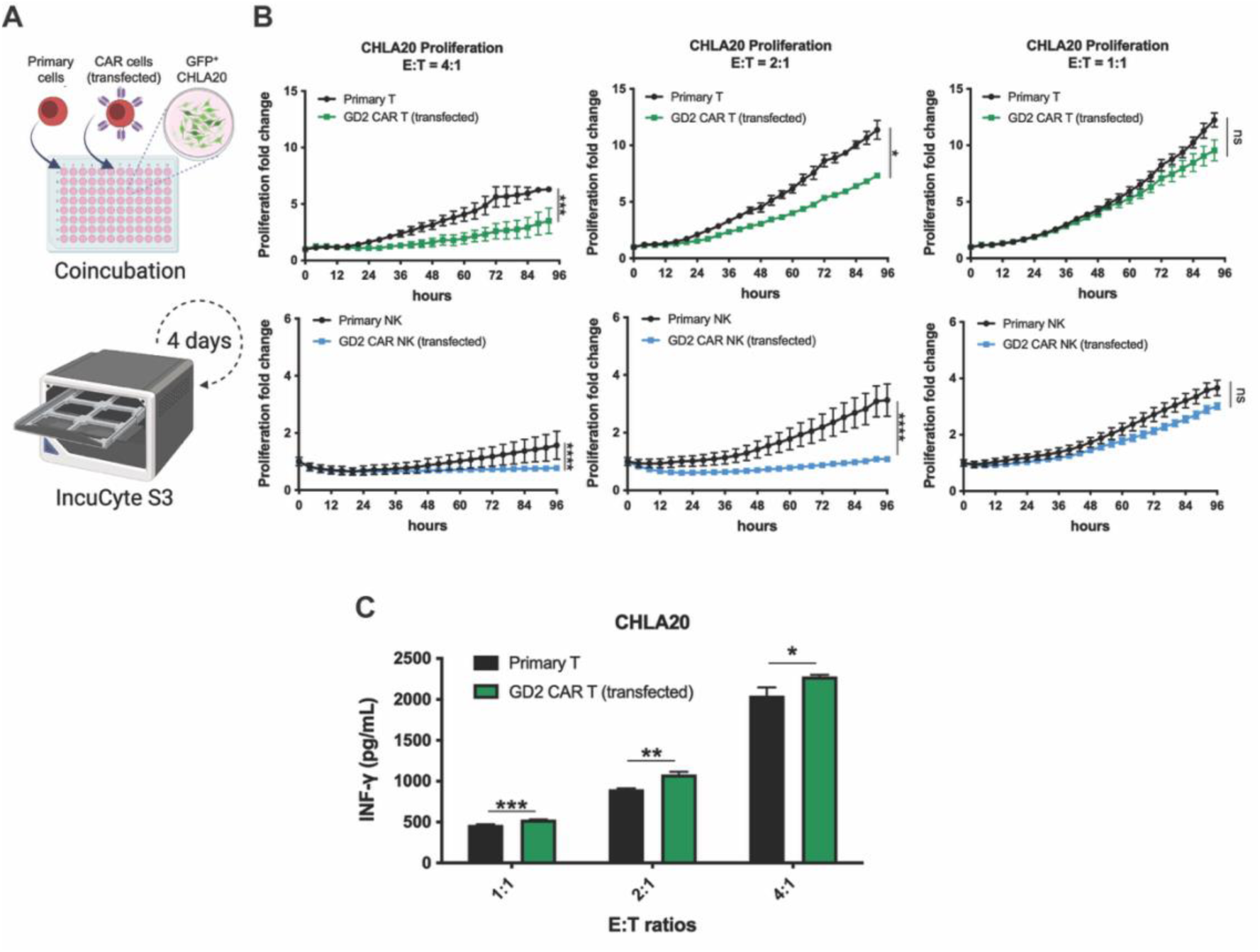
Potency of GD2 CAR T and CAR NK cells against neuroblastoma cells. (A) Schematic diagram of IncuCyte assay. (B) GFP^+^ CHLA20 cells were co-cultured with effector cells at various E:T ratios. The effector cells were either non-transfected (Primary) or CTC LNP-transfected (GD2 CAR). IncuCyte assay presented by proliferation fold change of CHLA20 cells recorded every 4 h. Data are presented as mean ± SEM (*n* = 3). (C) IFN-γ ELISA of GD2 CAR T cells and primary T cells 72 h after coincubation. Data are presented as mean ± SD (*n* = 3). ns, not significant; *p < 0.05; **p<0.01; ***p < 0.001; ****p < 0.0001, unpaired two-sided t-test. The scheme was created with BioRender.com

## Discussion

Since 2017, the FDA has approved seven CAR T cell products for treating hematological malignancies, including multiple myeloma, B-cell lymphomas, and B-cell acute lymphoblastic leukemia. Permanent CAR expression is achieved by all 7 products via viral transduction using retroviral or lentiviral vectors.^48,49^ However, this method raises several safety concerns that require additional regulatory compliance and monitoring, as well as significantly increased costs. Viral vectors can integrate into the host genome, potentially leading to insertional mutagenesis and oncogenic transformation.^48–50^ Even though replication-defective vectors are used, the FDA also requires monitoring for unintentional vector development of replication competence. Moreover, the use of viral transduction for CAR T cell production introduces high costs and complexity in manufacturing, in part due to the high cost of vector generation and bioprocessing at scale.^51^ Consequently, there is intense exploration of non-viral gene transfer methods to develop safer and more cost-effective CAR T cell therapies.^48^

In this study, we successfully developed an mRNA LNP utilizing FDA-approved lipids to generate both CAR T and CAR NK cells. A prior study demonstrated that an LNP containing ALC-0315 could effectively deliver mRNA to T cells *in vivo*.^52^ The ionizable lipid used in Pfizer-BioNTech’s tozinameran vaccine features an optimal alkyl tail length and a highly branched, symmetric structure, both of which are critical for efficient mRNA delivery.^53,54^ The helper lipid 1,2-dipalmitoyl-sn-glycero-3-phosphocholine (DPPC), along with cholesterol, provides structural stabilization and facilitates fusion with endosomal membranes.^55,56^ We selected the PEGylated lipid, 1,2-dimyristoyl-rac-glycero-3-methoxypolyethylene glycol-2000 (DMG-PEG2000), which has two C14 saturated alkyl chains that anchor its PEG moiety on the surface of LNPs. DMG-PEG2000 dissociates more rapidly from LNPs in serum-containing media compared to other PEGylated lipids with longer alkyl chains, making it ideal for *ex vivo* immune cell engineering that requires higher mRNA transfection efficiency.^57,58^ The molar ratios of these components and lipid selection can be optimized for specific applications, such as encapsulating various nucleic acid cargos, enhancing cellular uptake by target cells, altering biodistribution, and influencing protein corona formation.^59–64^ The versatility of LNPs enables the design of formulations tailored to the specific needs of mRNA-based CAR cell therapies, potentially improving their efficacy and safety profiles.

For reporter analysis, we observed the discrepancy in expression levels between the eGFP and mApple reporters. The results could be attributed to the 2A polycistronic construct, which favors higher expression of the first protein over the second protein in the mRNA transcript.^65^ Moreover, mApple has complex photoswitching behavior, a reversible bright-to-dark transition under wide-field illumination.^66,67^ The flickering effect may occur when multiple lasers are simultaneously illuminated in a flow cytometer, making mApple a less reliable reporter due to its inadequate emission intensity compared to eGFP.^68^ In the kinetic study, eGFP has a half-life in the order of days,^69^ which accounts for its sustained expression over an extended period observed in Jurkat cells, as well as in primary T and NK cells. Due to the photosensitivity issues associated with using mApple as a fluorescent reporter, we chose mCherry as the fluorescent reporter for our GD2 CAR mRNA transcript. Although fluorescent proteins may serve as surrogates in the kinetic study, further investigation is necessary to determine whether transfection with our LNP platform can sustain adequate CAR expression over time. Indeed, understanding the expression period of CAR is crucial for achieving the desired *in vivo* efficacy with mRNA-engineered cells. The information will aid in designing effective dosing regimens for CAR T and CAR NK cells produced through mRNA technology, ultimately leading to tumor growth suppression.^70–73^ The comparison between c-Myc-tagged CAR and mCherry expression in primary T cells revealed that, although the delivery of CAR mRNA was highly efficient, fewer cells were c-Myc-positive compared to those that were mCherry-positive. The observation may result from differences in protein complexity, translocation, and post-translational modifications between the two products.^74^ Moreover, the sensitivity of detecting GD2 CAR via the c-Myc tag, which relies on fluorophore-conjugated antibody staining in flow cytometry, may be lower than that of endogenous fluorescence from mCherry expressed in CTC LNP-transfected T cells.

The observation that exogenous human ApoE4 supplementation did not enhance mRNA delivery to primary T cells in serum-containing conditions indicates that fetal bovine serum (FBS) proteins can facilitate LNP uptake and cargo release. The approximately 80% mCherry positivity achieved with or without additional human ApoE4 suggests that the serum-containing medium already provides sufficient apolipoprotein isoforms of bovine origin to mediate effective transfection.^44^ This finding aligns with mechanistic studies demonstrating that upon LNP exposure to serum, rapid, sequential protein corona formation occurs, in which PEG-modified lipids are shed from the LNP surface within minutes, followed by dynamic rearrangement of the adsorbed protein layer to achieve an optimal uptake-competent state.^45,75^ This protein corona-mediated transformation reduces steric hindrance imposed by PEG coating and promotes lipid redistribution within the nanoparticle core, ultimately driving mRNA release into the cytoplasm. By proceeding with CTC LNP transfection experiments in serum-containing medium without supplemented human ApoE4, we leveraged the physiological complexity of endogenous FBS proteins to achieve efficient transfection while avoiding unnecessary experimental manipulation, demonstrating that the intrinsic composition of serum provides a naturally optimized vehicle for facilitating T cell uptake and mRNA delivery *in vitro*.

While CAR T cell therapies have been effective against hematological malignancies, their use as allogeneic cellular therapeutics is limited by their endogenous T-cell receptor (TCR), which can lead to alloreactive recognition of host antigens and an increased risk of graft-versus-host disease (GVHD).^76^ NK cells have emerged as a promising source for allogeneic CAR cell therapy since they do not express TCR and cannot mediate GVHD. Thus, CAR NK cells offer great potential to become off-the-shelf products, resulting in lower manufacturing costs and reduced treatment delays compared to autologous immune cell therapies.^77^ Furthermore, CAR NK cells have the innate ability to engage stress ligands in conjunction with engineered CARs, which may enhance antitumor responses.^78^ Ongoing clinical trials are exploring the applications of CAR NK cells in both hematological and solid tumors.^79^ Although viral transduction can achieve stable CAR expression, NK cells exhibit significantly lower transduction efficiency compared to T cells due to their robust innate antiviral defenses. Pattern recognition receptors (PRRs), such as Toll-like receptors (TLRs) and cytosolic nucleic acid sensors, can detect foreign genetic material introduced during viral vector-based transduction.^80,81^ Upon recognition of foreign genetic materials, receptors trigger intracellular signaling cascades that activate pro-apoptotic pathways and inflammatory responses, contributing to the poor survival of transduced NK cells.^82^ As a result, researchers are investigating alternative approaches, such as lipid-based mRNA delivery systems, for more efficient and viable NK cell engineering.

The introduction of CAR constructs to primary T and NK cells through downstream *ex vivo* modifications is influenced by factors such as the cellular differentiation state, degree of exhaustion, and proliferative capacity.^83–85^ T and NK cells present unique challenges for mRNA delivery systems due to their inherent resistance to particle uptake, necessitating proper activation before transfection. Robust activation and expansion of primary T cells are typically achieved through co-stimulation with CD3 and CD28, which is essential for effective LNP-based transfection.^10,12,86^ Our results supported that primary NK cells successfully activated by K562-mbIL15-41BBL feeder cells exhibited high transfection efficiencies, highlighting the critical relationship between cell proliferation capacity and subsequent cell engineering. Successful transfection of the LNP platform likely requires an activation process that increases metabolic activity, improves endocytosis, and facilitates the translation machinery in primary T and NK cells.

Further research into the mechanisms underlying the phenotypic shifts of immune cells following LNP treatment could provide valuable insights for developing next-generation LNPs for immune cell engineering. Our analysis of NK cells revealed significant phenotypic changes due to LNP treatment, including reduced NKp46 expression and a shift from CD16^+^CD56^bright^ toward CD16^dim^CD56^bright^ NK cell populations. The changes were observed irrespective of whether the LNPs carried mRNA cargo, suggesting that the lipid components alone can influence NK cell phenotype. Such alterations may have functional implications: NKp46 is a primary activating receptor involved in tumor recognition^87,88^, while a shift in CD16/CD56 expression could affect antibody-dependent cellular cytotoxicity and cytokine production.^47,89,90^ Although transfection of NK cells using CTC LNPs yields lower mCherry reporter expression than T cells, we can further optimize the full potential of mRNA LNP-engineered NK cells by refining LNP formulations and mRNA designs to enhance CAR protein expression and better preserve desired NK cell phenotypes.

In conclusion, we developed a non-viral method for generating GD2 CAR T and CAR NK cells using mRNA-loaded LNPs. Our LNP formulation, composed of lipids used in FDA-approved products, demonstrated high transfection efficiency, minimal cytotoxicity, and the ability to produce functional GD2 CAR cells that show potency against GD2^+^ neuroblastoma cells. The success of this method in engineering T and NK cells addresses several limitations of current viral vector-based approaches, including safety concerns and manufacturing complexities.

## Materials and methods

### Cell culture

Jurkat human T cell lymphoma cells were generously provided by Dr. Quanyin Hu at the University of Wisconsin-Madison. NK-92 human NK cell lymphoma cells were obtained from ATCC (#CRL-2407, Manassas, VA). The AkaLUC-GFP CHLA20 human neuroblastoma cells were a gift from Dr. James Thomson at the Morgridge Institute for Research and maintained in Dulbecco’s Modified Eagle Medium (DMEM) with 4.5 g/L glucose, L-glutamine, and sodium pyruvate (Corning). Jurkat and primary human T cells were cultured in RPMI-1640 with L-glutamine (Corning, Corning, NY). K562-mb15-41BBL cells were a gift from Dr. Dario Campana (National University of Singapore) and maintained in RPMI-1640 with L-glutamine. Mouse T cells were cultured in R10 medium, which was prepared using RPMI-1640 with L-glutamine, supplemented with 50 mM HEPES (Corning), 1 mM sodium pyruvate (Corning), and 55 nM 2-mercaptoethanol (Thermo Fisher Scientific, Waltham, MA). Primary human NK cells were cultured in RPMI-1640 with L-glutamine (Corning) by including an additional 10 mM HEPES (Corning), 1 mM sodium pyruvate (Corning), and 1X MEM Nonessential Amino Acids (Corning). CHLA20 cells were maintained in Dulbecco’s Modified Eagle Medium (DMEM) with 4.5 g/L glucose, L-glutamine, and sodium pyruvate (Corning). NK-92 were cultured in DMEM supplemented with 10 µM 2-mercaptoethanol (Thermo Fisher Scientific). All of the above culture media were supplemented with 10% fetal bovine serum (FBS) (MilliporeSigma, Burlington, MA) and 1% penicillin-streptomycin (Corning). ATCC guidelines were followed for cell authentication using morphology monitoring, growth curve analysis, and testing for mycoplasma within 6 months of use. All cells were cultured in an incubator (Galaxy 170 S, Eppendorf, Germany) at 37℃ with an atmosphere of 5% CO2.

### mRNA synthesis

The plasmid DNA encoding pT7-eGFP-T2A-mApple (VectorBuilder, Chicago, IL) was linearized by SmaI restriction enzymes (New England Biolabs, Ipswich, MA), giving the DNA template for *in vitro* transcription (IVT). The eGFP-T2A-mApple mRNA was generated via IVT using a HiScribe T7 High Yield RNA Synthesis Kit (New England Biolabs) with co-transcriptional 5’ capping using the CleanCap® Reagent AG (3’ OMe) (TriLink Biotechnologies, San Diego, CA). UTPs were fully substituted with pseudo-UTP (TriLink Biotechnologies) in the IVT reaction, which was incubated for 2 h at 37℃. The resultant mRNA was treated with 100 IU/mL DNase I (New England Biolabs) for 15 min at 37℃ to eliminate the DNA template. Subsequent polyadenylation was performed for 30 min at 37℃ using *E. coli* Poly(A) Polymerase (New England Biolabs). The mRNA was purified using the Monarch® RNA Cleanup Kit (New England Biolabs), followed by verification of the concentration using a NanoQuant Plate (Tecan, Männedorf, Switzerland). The purified mRNA was then stored at −80℃ until use in LNP formulation. The pT7-SP-14G2A-T2A-mCherry (VectorBuilder) plasmid, which already contained the poly(A) sequence, was linearized by HindIII restriction enzyme (New England Biolabs). The GD2 CAR sequence 14G2a-41BB-CD3ζ was a gift from Dr. Crystal Mackall at Stanford University, and GD2 CAR-T2A-mCherry mRNA was generated via the same IVT and purification procedures as the reporter constructs.

### LNP formulation and characterization

All LNPs were formulated using a pipette mixing method as described previously.^34^ Briefly, an organic phase was prepared by dissolving ionizable lipid (ALC-0315) (Avanti Polar Lipids, Birmingham, AL), helper lipid (DPPC) (Avanti Polar Lipids), cholesterol (MilliporeSigma), and PEG-lipid (DMG-PEG2000) (Avanti Polar Lipids) in ethanol to a final concentration of 1 mM with the molar ratios of 50:10:38.5:1.5, respectively.^91^ The ionizable lipid nitrogen to mRNA phosphate (N/P) ratio was fixed at 6. An aqueous phase containing mRNA was prepared using 50 mM citrate buffer at pH 3.5. Organic and aqueous phases were mixed at a 1:3 volume ratio via rapid pipetting for 30 sec to generate LNPs. LNPs were then dialyzed against 1× Dulbecco’s Phosphate-Buffered Salt Solution (DPBS) (Corning) using a Slide-A-Lyzer Dialysis Cassette, 10K MWCO (Thermo Fisher Scientific) for 24 h under 4℃, concentrated at 1,300 𝑔 for 10 min under 4℃ using an Amicon® Ultra Centrifugal Filter, 30 kDa MWCO (MilliporeSigma). The preparation of blank LNPs followed the same procedure, but the aqueous phase contained no mRNA molecules. LNPs were characterized using a Zetasizer Nano ZS instrument (Malvern Panalytical, Malvern, UK). Briefly, 100 μL of LNP suspension was diluted in 600 μL of Milli-Q water in a disposable cuvette (Malvern Panalytical) suitable for measuring zeta potential, size, and polydispersity.

### LNP encapsulation

LNP encapsulation efficiency was determined using a Quant-iT RiboGreen assay kit (Thermo Fisher Scientific) as previously described.^92^ Two RNA standard curves were prepared via serial dilution in 1:1 DPBS and 2× TE supplemented with or without 1% Triton X-100. For external mRNA measurement, LNPs collected after the centrifugal filtration were diluted 2-fold using 2× Tris-EDTA (TE) buffer. For total mRNA measurement, LNPs were first diluted 10-fold in DPBS, then diluted 2-fold in 2× TE buffer containing 1% (v/v) Triton X-100 (MilliporeSigma), and incubated for 20 min to lyse the particles. The RiboGreen solution in 1X TE was mixed 1:1 with the standards or LNP samples in a black 96-well plate (Thermo Fisher Scientific) according to the manufacturer’s instructions. Fluorescence intensity was measured using a CLARIOstar Plus microplate reader (BMG Labtech, Ortenberg, Germany) at 480 nm (excitation) and 520 nm (emission). The standard curve with or without Triton X-100 was used to quantify total mRNA and external mRNA, respectively. Relative encapsulation efficiency is calculated as 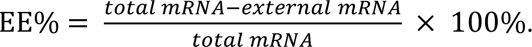

### LNP pKa and surface ionization

LNP apparent p*K*a was identified as previously described.^93^ Briefly, a buffer solution consisting of 150 mM sodium chloride, 20 mM sodium phosphate, 20 mM ammonium acetate, and 25 mM ammonium citrate was titrated to pH values ranging from 2 to 12 using 3 M hydrochloric acid and 5 M sodium hydroxide. A black 96-well plate (Thermo Fisher Scientific) was loaded with 250 μL of each pH solution in triplicate, and 5 μL of LNP was added to each well. A 0.16 mM 6-(p-Toluidino)-2-naphthalenesulfonic acid (TNS) solution was prepared as a stock, and 10 μL of the TNS solution was added to each well, resulting in a final concentration of 6 μM. Fluorescence intensity was measured using a CLARIOstar Plus microplate reader (BMG Labtech) at an excitation wavelength of 322 nm and an emission wavelength of 431 nm. The TNS assay data were interpolated with an asymmetric sigmoidal curve and analyzed with Graphpad Prism (Dotmatics, Boston, MA). The apparent p*K*a of LNP was determined as the pH corresponding to half-maximum fluorescence intensity, representing 50% protonation of the LNP surface.

### Cell viability

The Jurkat cell line was used to perform the cell viability assay. Cells were cultured in RPMI-1640 containing 10% FBS and 1% penicillin-streptomycin. Cells were plated at 10,000 cells per well in a 96-well plate (Corning) in 100 μL of media. After 24 h of incubation at 37℃, cells were treated with the blank LNP at 0.09, 0.18, 0.36, 1.08, 2.16, 10.8, and 21.6 nmol of total lipids per 10,000 cells or the mRNA-loaded LNP at 2.5, 5, 10, 30, 60, 300, and 600 ng of mRNA per 10,000 cells, and allowed to incubate at 37℃ for 24 h. The total lipids of the mRNA-loaded LNP matched the blank LNP at each dose. After the treatment, 10 μL of CellTiter-Blue® Cell Viability Assay (Promega, Madison, WI) was added to each well and incubated with the cells for an additional 3 h at 37℃. The fluorescence intensity, measured with excitation at 560 nm and emission at 590 nm, was analyzed using a Synergy H1 plate reader (BioTek, Winooski, Vermont). The cell viability was calculated as a percentage of the untreated control.

### In vitro mRNA delivery to Jurkat and NK-92 cells

Jurkat cells were plated at 1 × 10^5^ cells/mL, and NK-92 cells were plated at 2 × 10^5^ cells/mL in 24-well plates (Corning). Jurkat and NK-92 cells were treated with 300 ng of LNP-encapsulated mRNA per 10,000 cells. For the LNPs, the amount of mRNA was calculated based on the formulation volume obtained after dialysis and centrifugation. One day after transfection, fluorescent images were captured on an Axio Vert.A1 microscope (Zeiss, Oberkochen, Germany) using a 20× objective, followed by flow cytometry experiments. The fluorescent reporters expressed by Jurkat cells were observed daily for 2 weeks using the fluorescent microscope. During the kinetic study of protein expression, the cells were supplemented with the culture medium every other day to maintain cell growth. All images were processed with Fiji software version 2.16 for background subtraction, contrast adjustment, and brightness enhancement.

### In vitro mRNA delivery of murine T cells

Eight- to ten-week-old female C57BL/6NCr (B6) mice (Charles River Laboratories, Wilmington, MA) were used in this study. All mice were housed and cared for in accordance with the Guide for the Care and Use of Laboratory Animals. All animal experiments were approved by the Institutional Animal Care and Use Committee (IACUC) under protocol M005647. To isolate murine T cells for *in vitro* experiments, spleens were excised from B6 mice and strained through a 100-μm filter into R10 medium. Splenocytes were collected via centrifugation. After removing the supernatant, the cell pellet per spleen was resuspended in 5 mL of ACK Lysing Buffer (Lonza, Walkersville, MD) and incubated on ice for 5 min to lyse red blood cells. An equal volume of R10 media was added, followed by centrifugation and resuspension of the splenocytes. The splenocytes were used to isolate murine CD3^+^ T cells using the MojoSort Mouse CD3 T Cell Isolation Kit (BioLegend, San Diego, CA) following the manufacturer’s protocol. Isolated murine T cells were cultured overnight at 1 × 10^6^ cells/mL in R10 media containing 30 IU/mL rhIL-2. The cells were then activated for 24 h using Dynabeads Mouse T-Activator CD3/CD28 (Thermo Fisher Scientific) with a 1:1 bead-to-cell ratio. After activation, cells were treated with LNPs at 30 or 60 ng of encapsulated mRNA per 10,000 cells. Murine T cells expressing fluorescent reporters were observed under a 20× objective on an Axio Vert.A1 microscope and measured by flow cytometry 24 h after transfection with LNPs.

### Ex vivo mRNA delivery to primary human T cells

Human CD3^+^ T cells were isolated from human PBMCs obtained from healthy volunteers (IRB protocol 2019-0865) using the MojoSort Human CD3 T Cell Isolation Kit (BioLegend) following the manufacturer’s protocol. Human CD3^+^ T cells were plated at 1 × 10^6^ cells per well in 1 mL of RPMI-1640 in 24-well plates precoated with 1.5 μg anti-CD3 antibody (Thermo Fisher Scientific) at 37℃ for 2 hours. The cells were activated for 72 h with 1 μg/mL of anti-CD28 (Thermo Fisher Scientific) and 150 IU/mL rhIL-2 (Hoffmann-La Roche Inc., Nutley, NJ) supplemented in the culture medium. After activation, cells were plated at 1 × 10^6^ cells/mL and treated with LNPs at 30 or 60 ng of encapsulated mRNA per 10,000 cells. Human T cells expressing fluorescent reporters were observed using a 20× objective on an Axio Vert. A1, and measured via flow cytometry one day after transfection with LNPs. The kinetics of the fluorescent reporters were monitored daily for 8 days. The cells were supplemented with the same culture medium containing 150 IU/mL rhIL-2 every other day and split into two wells to maintain cell growth during the expression kinetic study. All images were processed with Fiji software version 2.16 for background subtraction, contrast adjustment, and brightness enhancement. For GD2 CAR mRNA experiments, the same *ex vivo* transfection protocol was used. To assess the dependency of GD2 CAR mRNA delivery on human ApoE4-mediated endocytosis, T cells were transfected with mRNA LNPs in the absence or presence of 1 μg/mL of human ApoE4 (PeproTech, Cranbury, NJ) supplemented in serum-containing medium. The generation of GD2 CAR T cells was first examined by observing mCherry using an Axio Vert.A1 microscope under a 20× objective, followed by analyzing the expression of the mCherry reporter and c-Myc-tagged GD2 CAR via flow cytometry.

### Primary NK cell expansion

NK cells were isolated from peripheral blood mononuclear cells (PBMCs) obtained from healthy donors (IRB protocol 2017-1070) using the RosetteSep™ Human NK Cell Enrichment Cocktail (STEMCELL Technologies, Vancouver, Canada). Isolated NK cells were co-cultured with irradiated (100 Gy) K562-mbIL15-41BBL feeder cells at a feeder-to-NK cell ratio of 2:1. The cells were maintained in RPMI 1640 medium supplemented with 10% FBS, 1X MEM non-essential amino acids (Corning), 1% penicillin/streptomycin, 100 µM β-mercaptoethanol (Thermo Fisher Scientific), and 100 IU/mL rhIL-2 (NCI, Bethesda, MD). Culture media were refreshed every 2–3 days. Total NK cells were counted 3, 6, and 8 days after feeder cell activation using Trypan Blue Solution (Corning).

### Cell cycle analysis

Half a million to one million activated NK cells were harvested 3, 6, and 8 days after feeder cell expansion and resuspended in 300 μL of PBS. Cold ethanol (700 μL) was added dropwise to a final concentration of 70%. Samples were fixed at 4℃ overnight and then centrifuged at 500 𝑔 for 5 min and washed with 1 mL PBS twice. The cells were resuspended with 500 μL of FxCycle PI/RNase Staining Solution (Thermo Fisher Scientific). The samples were wrapped in foil to protect from light and incubated for 30 min at RT. Cell cycle data were acquired on an Attune NxT flow cytometer (Thermo Fisher Scientific) and analyzed using FCS Express 7 software (Dotmatics).

### Ex vivo mRNA delivery to primary human NK cells

LNP transfection of primary NK cells was performed after 8 days of feeder cell activation. After feeder cell activation, expanded human NK cells were plated at 2 × 10^6^ cells/mL in 48-well plates (Greiner Bio-one, Frickenhausen, Germany) and treated with LNPs at 30 ng of encapsulated eGFP-T2A-mApple mRNA per 10,000 cells. Human NK cells expressing eGFP and mApple fluorescent reporters were measured via flow cytometry one day after transfection with LNPs, and the kinetics of the fluorescent reporters were monitored for one week. The cells were supplemented with fresh culture medium containing 100 IU/mL rhIL-2 every other day.

For GD2 CAR mRNA delivery, the same NK expansion and transfection protocol was used. The generation of GD2 CAR NK cells was verified by mCherry reporter expression using flow cytometry.

### Flow cytometry

Flow cytometry was performed on an Attune NxT flow cytometer (Thermo Fisher Scientific) with blue, violet, yellow, and red lasers. All experiments used Ghost Dye Red 780 (Cytek Biosciences, Fremont, CA) as a viability stain. Briefly, cells in 1 mL DPBS were incubated with 1 μL of the dye at 4℃ for 30 min, then quenched with 1 mL of FACS buffer (PBS, 5% FBS and 0.2% sodium azide). Compensation matrices were obtained using singly stained OneComp eBeads (Thermo Fisher Scientific) or stained cells. For c-Myc-tagged CAR staining, T cells were incubated with 1 μL of c-Myc biotinylated antibody (Miltenyi Biotec, Bergisch Gladbach, Germany) at 4℃ for 30 min followed by 1 μL of anti-biotin PE (Miltenyi Biotec) for another 30 min. The primary NK cells were stained with 1 μL of antibodies at 4℃ for 30 min, including anti-CD56-AF647 and anti-CD3-BV605 (BioLegend) for gating the NK population. For the phenotype study, non-treated, blank LNP-treated, and mRNA LNP-treated NK cells were stained with 1 μL of antibodies at 4℃ for 30 min, including anti-CD56-AF647, anti-CD3-BV605, anti-CD16-BV421, and anti-NKp46-PE-Cy7 (BioLegend). Flow cytometry data were analyzed using FCS Express software (Dotmatics).

### IncuCyte assays

AkaLUC-GFP CHLA20 cells were seeded at 10,000 cells per well in a 96-well plate. After 24 hours, CTC LNP-transfected and non-transfected cells were added to wells at 4:1, 2:1, and 1:1 effector-to-target (E:T) ratios in triplicate. The plate was centrifuged at 300 𝑔 for 1 min and then placed in the IncuCyte® S3 Live-Cell Analysis System (Sartorius, Göttingen, Germany), stored inside a cell incubator at 37℃ and 5% CO2. Images were taken every 4 h for approximately 4 days, and the proliferation of GFP^+^ CHLA20 cells was calculated with the following formula: proliferation [x-fold] = GFP^+^ cells at image time/initial GFP^+^ cells.

### ELISA assays

ELISA was performed to analyze the supernatant collected from the IncuCyte assay of GD2 CAR T cells co-cultured with GD2^+^ AkaLUC-GFP CHLA20 cells. According to the manufacturer’s protocol, human interferon-gamma (IFN-γ) levels were examined using the IFN-γ ELISA Kit (BioLegend). The absorbance intensity at 450 nm was read using an Epoch Microplate Spectrophotometer (BioTek), and the concentration of IFN-γ was calculated using a standard curve.

### Statistics

Statistical data and graphs were generated using GraphPad Prism 10 software (Dotmatics). A two-sided unpaired Mann-Whitney (nonparametric data) or t-test (parametric data) was used to determine the significance of differences between two groups. A two-way ANOVA with Dunnett’s or Sidak’s multiple comparison test was used to compare the means of 3 or more groups simultaneously. Statistical significance was defined as a p-value less than 0.05.

## Data availability

Data supporting the findings of this study are available upon reasonable request.

## Acknowledgments

Funding for this work was provided by NIH/NCI R01 CA278051 (CMC), St. Baldrick’s Foundation Empowering Immunotherapies for Childhood Cancer (EPICC) research grant (CMC), and the Midwest Athletes Against Childhood Cancer (MACC) Fund (CMC and JG). A donation from the Lachman Institute for Pharmaceutical Development to SM supported the initial development of the lipid nanoparticles and their physicochemical characterization.

We also acknowledge the University of Wisconsin Carbone Cancer Center Flow Cytometry Laboratory supported by NCI P30 CA014520. The contents of this article do not necessarily reflect the views or policies of the Department of Health and Human Services, nor does mention of trade names, commercial products, or organizations imply endorsement by the US Government.

## Author contributions

CC drafted the manuscript, designed LNPs, conducted experiments, performed statistical analysis, and analyzed data; LS and SHC designed the DNA constructs for mRNA IVT; LS optimized NK cell activation and expansion; SHC supervised CC on mRNA IVT and synthesized mRNA; AP supervised CC on flow cytometry and data analysis; VV contributed to mRNA LNP preparation and characterization. SM interpreted data and supervised CC and VV on LNP development; CMC interpreted data and supervised CC and LS on NK cell engineering; JG interpreted data and supervised CC on experimental design. JG conceived the research project with SM and CMC. All authors have revised and approved the manuscript.

