## Supplemental Materials for "Synthesis of mRNA lipid nanoparticles for engineering GD2 CAR T and CAR NK cells against neuroblastoma"

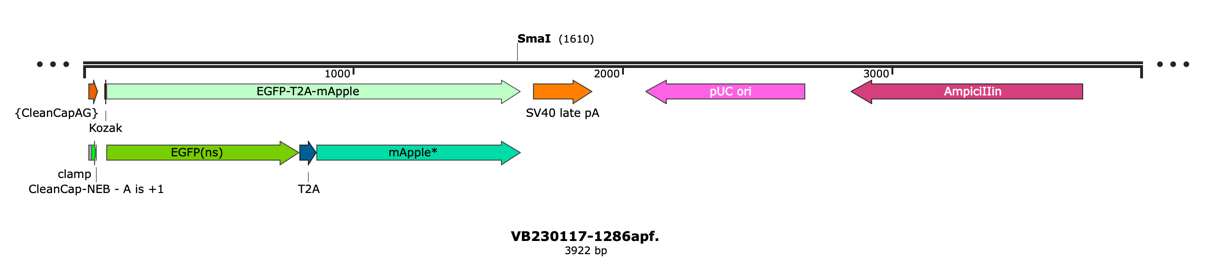

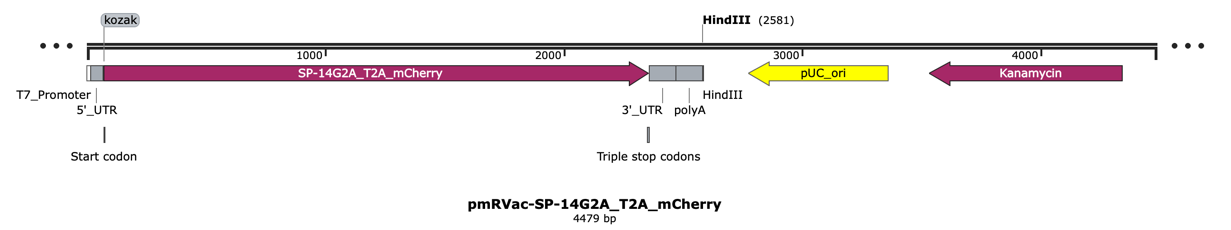

**Figure S1. DNA templates for mRNA IVT.** The top DNA construct is eGFP-T2A-mApple, which encodes a green fluorescent protein (eGFP) and a red fluorescent protein (mApple). This mRNA (~1,650 bases) was used for LNP optimization and validation. The bottom construct is GD2 CAR-T2A-mCherry, which contains the anti-GD2 CAR and mCherry sequences (~2,400 bases). Both plasmids were designed for mRNA IVT using T7 polymerase. The DNA template allowed co-transcriptional 5'-capping using CleanCap® Reagent AG. Specific cutting sites of the restriction enzymes were included for linearizing the plasmid. The 2A self-cleaving peptide, T2A, was inserted between the protein sequences of both constructs to induce ribosomal skipping during translation, thereby allowing the generation of two different proteins from a single mRNA transcript. The DNA maps were created with SnapGene.

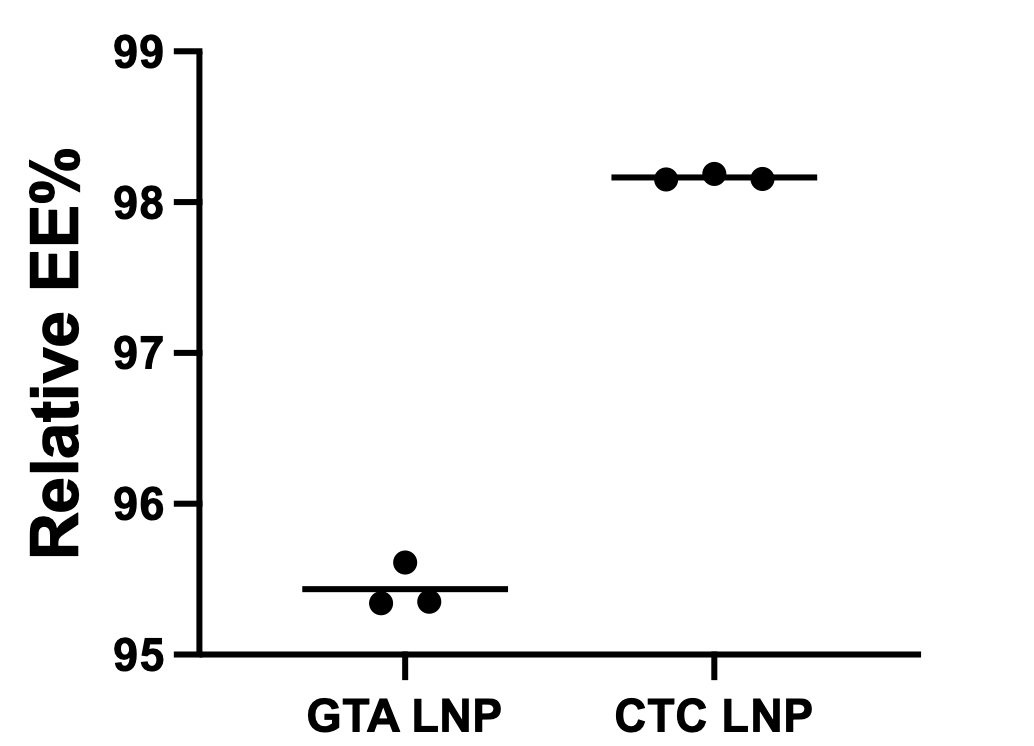

**Figure S2. Relative encapsulation efficiency of GTA and CTC LNPs.** The encapsulation efficiency (EE%) was determined for LNPs prepared with GTA or CTC mRNA. Data are presented as technical replicates (*n* = 3), with the horizontal line representing the mean.

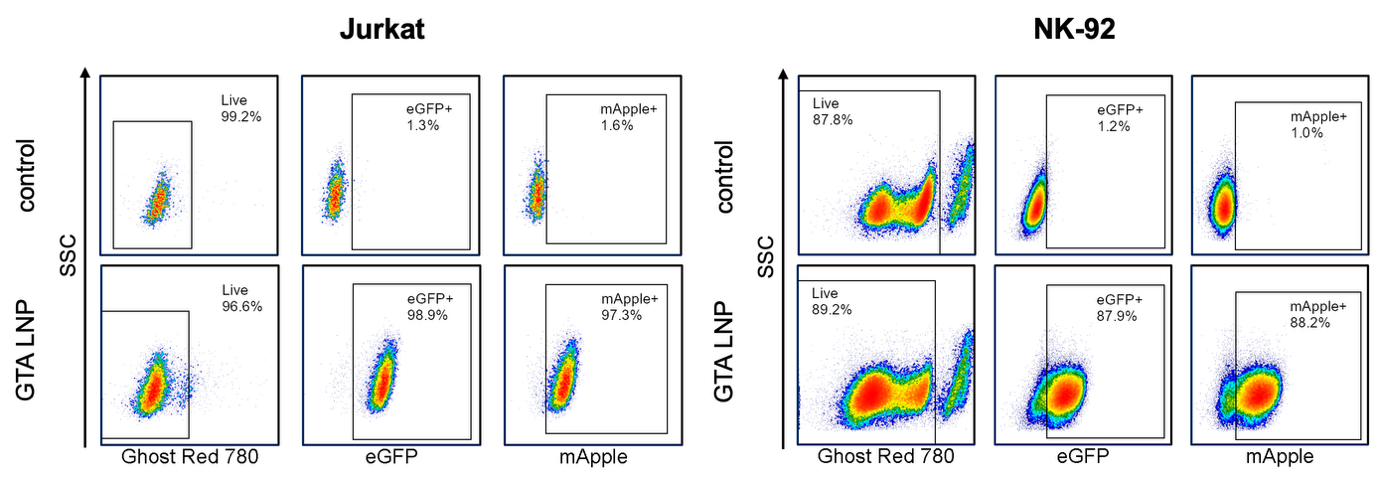

**Figure S3.** **GTA mRNA delivery into Jurkat and NK-92 cells by LNPs.** Representative flow cytometric density plot of Ghost Dye Red staining and fluorescent reporter expression of Jurkat and NK-92 cells.

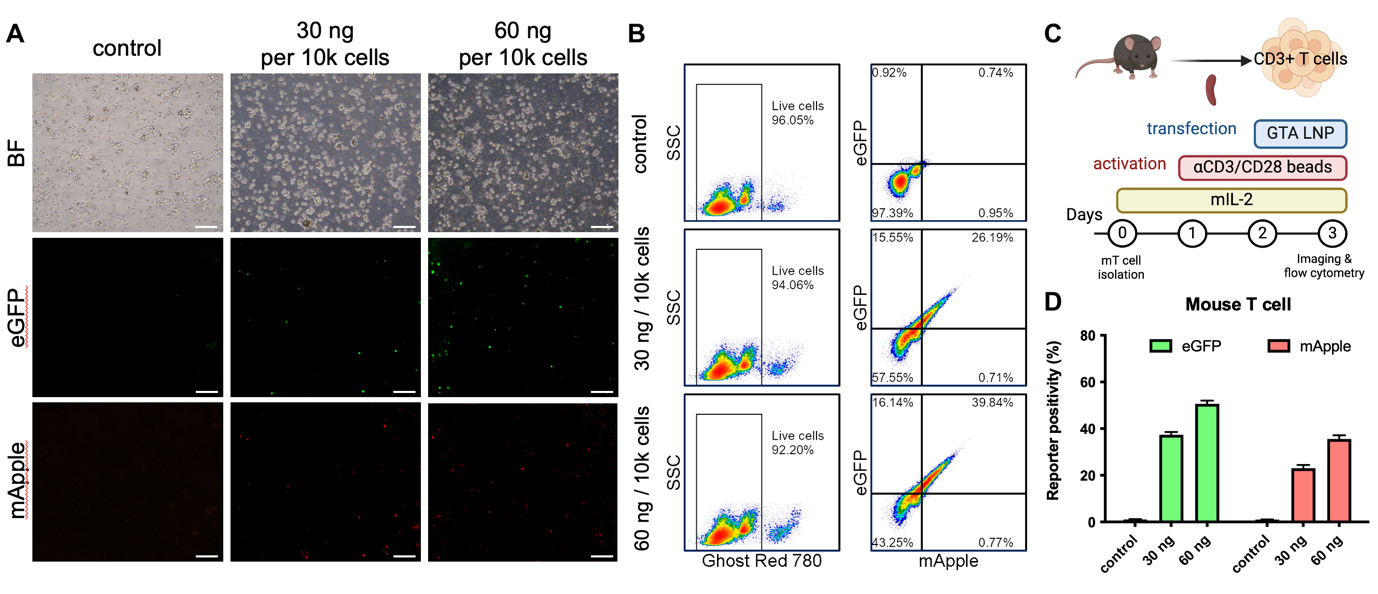

**Figure S4. GTA mRNA delivery into murine T cells by LNPs.** (A) Representative bright-field (BF) and fluorescent images of murine T cells with or without LNP transfection. Scale bar = 100 μm. (B) Representative flow cytometric density plot of Ghost Dye Red staining and fluorescent reporter expression of the mouse T cells. (C) Experimental design for *in vitro* murine T cell transfection. Murine T cells were magnetically isolated from murine splenocytes and supplemented with 30 IU/mL rhIL-2 every other day. After overnight incubation, the cells were activated for 24 hours with αCD3/CD28 beads before LNP transfection. (D) Reporter positivity of the LNP-transfected mouse T cells was analyzed via flow cytometry. Data are presented as mean ± standard deviation (*n* = 3). The scheme was created with BioRender.com.

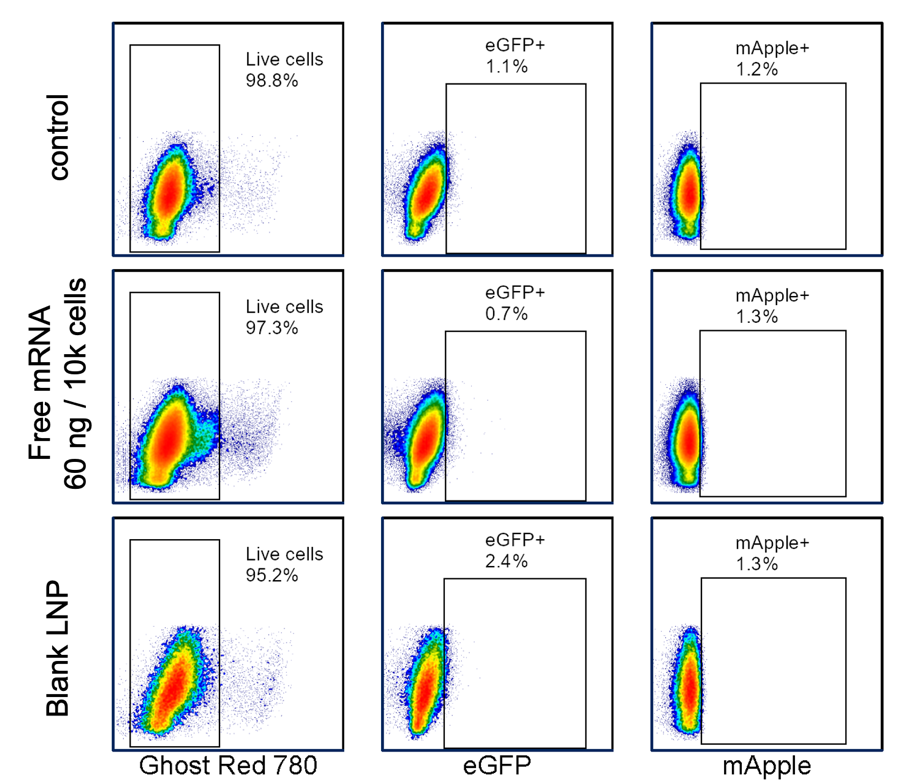

**Figure S5. Free mRNA and blank LNP transfection on primary human T cells.** Representative flow cytometric density plot of live-dead staining and reporter expression of untreated control, primary human T cells treated with free mRNA, or blank LNP.

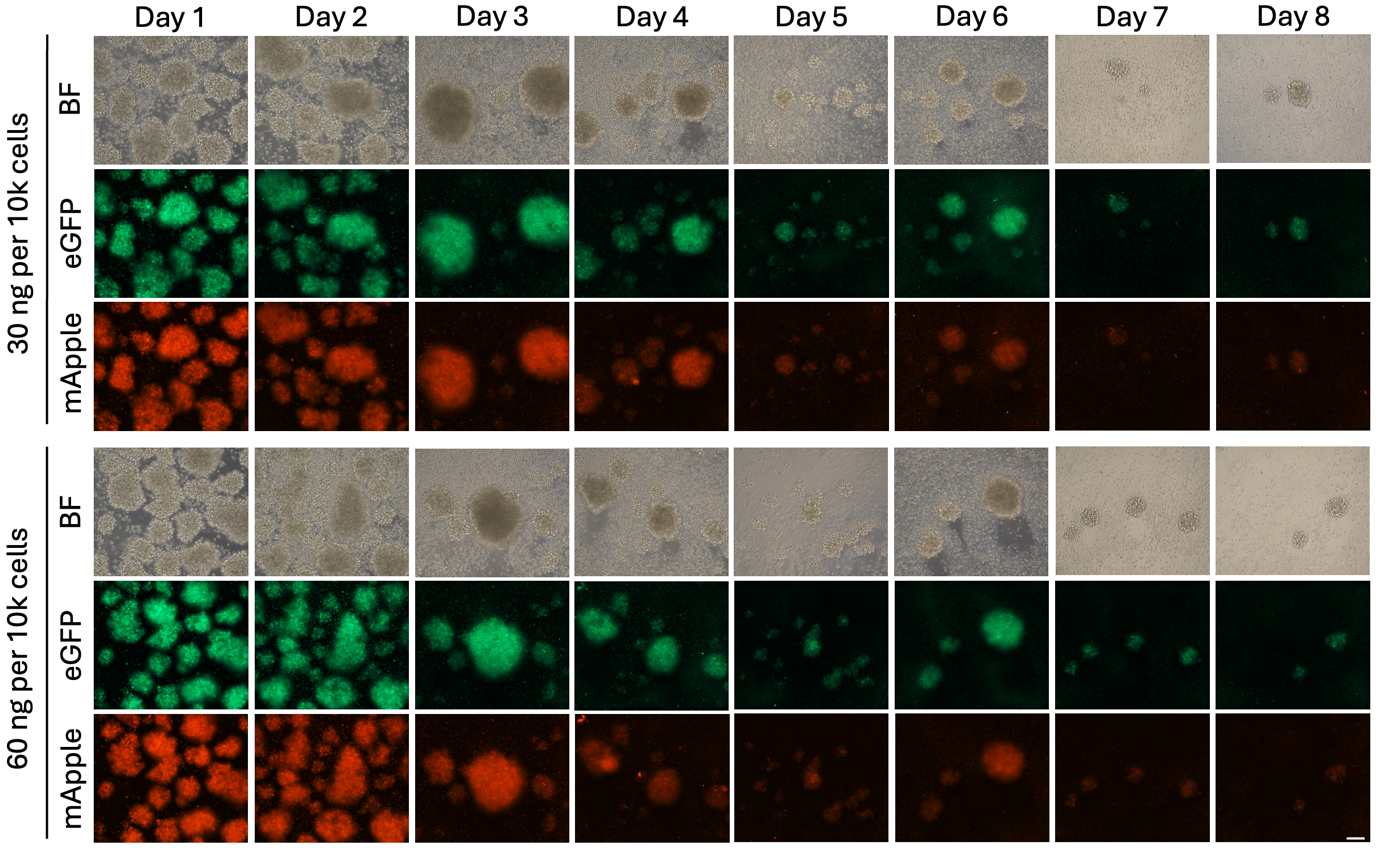

**Figure S6. Fluorescent proteins expressed by *ex vivo* engineered primary human T cells.** Representative fluorescent images showing eGFP and mApple expression of primary human T cells from 1 to 8 days post-transfection with 30 or 60ng LNP per 10,000 (10k) T cells. Scale bar = 100 μm. BF= brightfield.

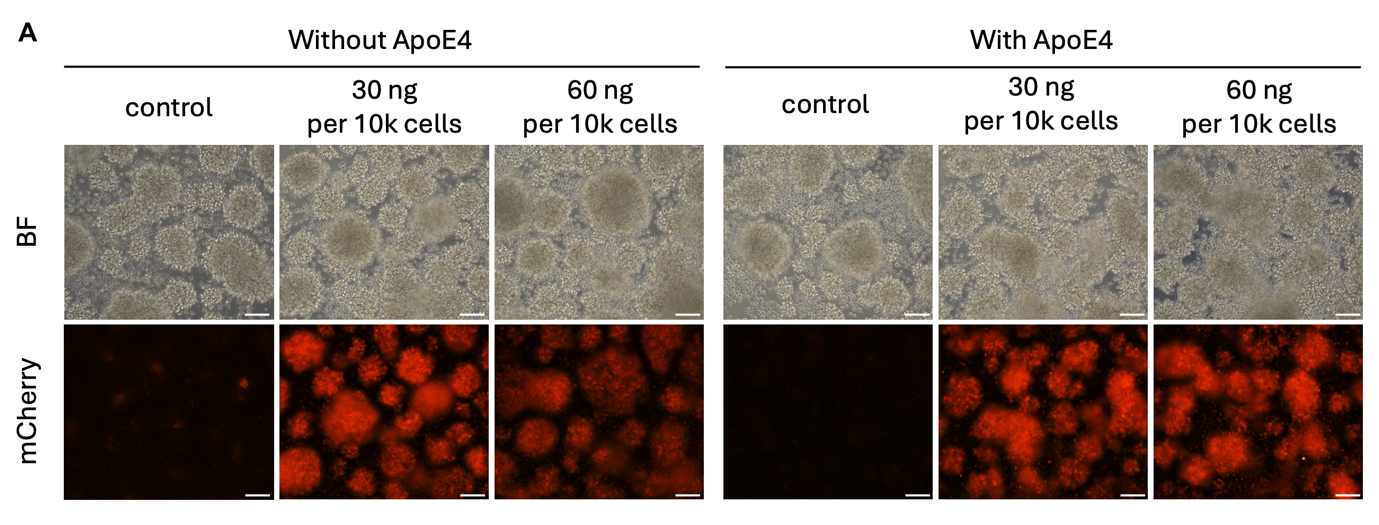

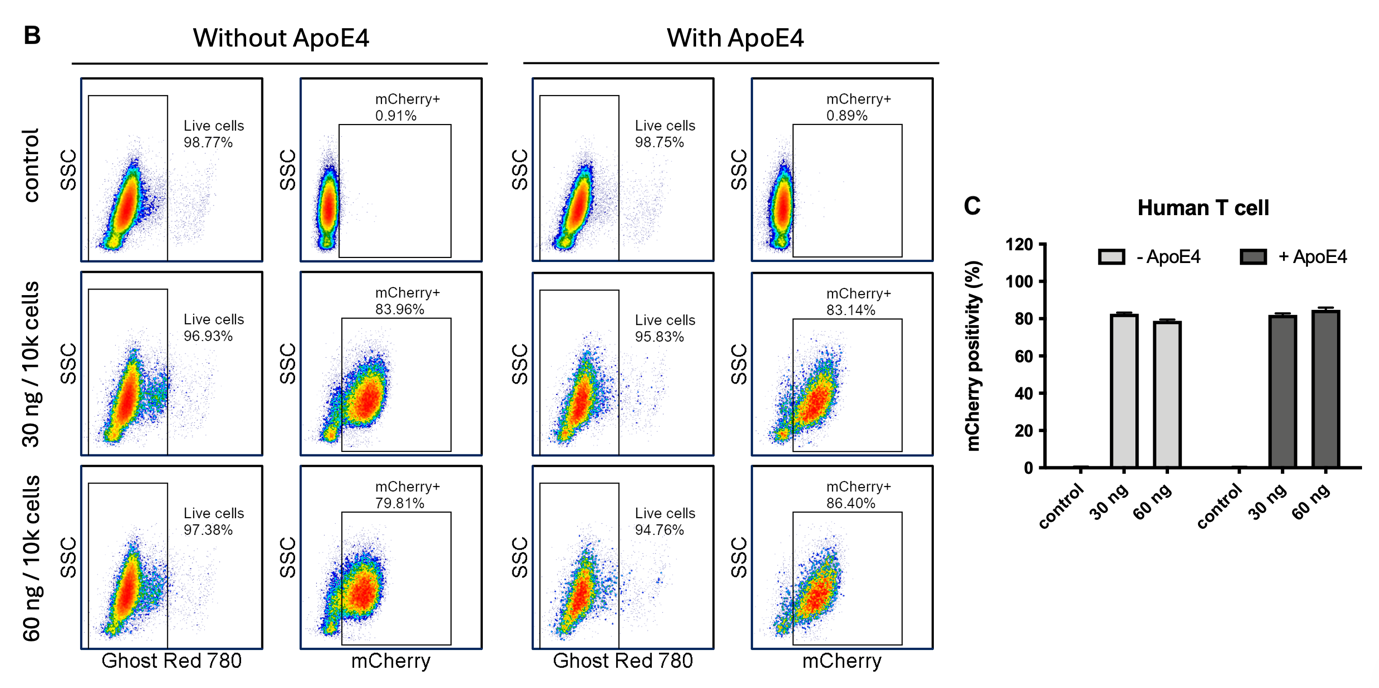

**Figure S7. Effect of additional human ApoE4 on primary human T cell transfection.** Primary human T cells were transfected with CTC LNP containing the corresponding doses of GD2 CAR-T2A-mCherry mRNA for 24 h with or without 1 μg/mL of human ApoE4. (A) Representative brightfield (BF) and mCherry images of the transfected T cells. Scale bar = 100 μm. (B) Representative flow cytometric density plot of viability staining and mCherry expression of primary human T cells. (C) mCherry positivity of primary human T cells transfected at low and high doses of LNP. Data are presented as mean ± standard deviation (*n* = 3).

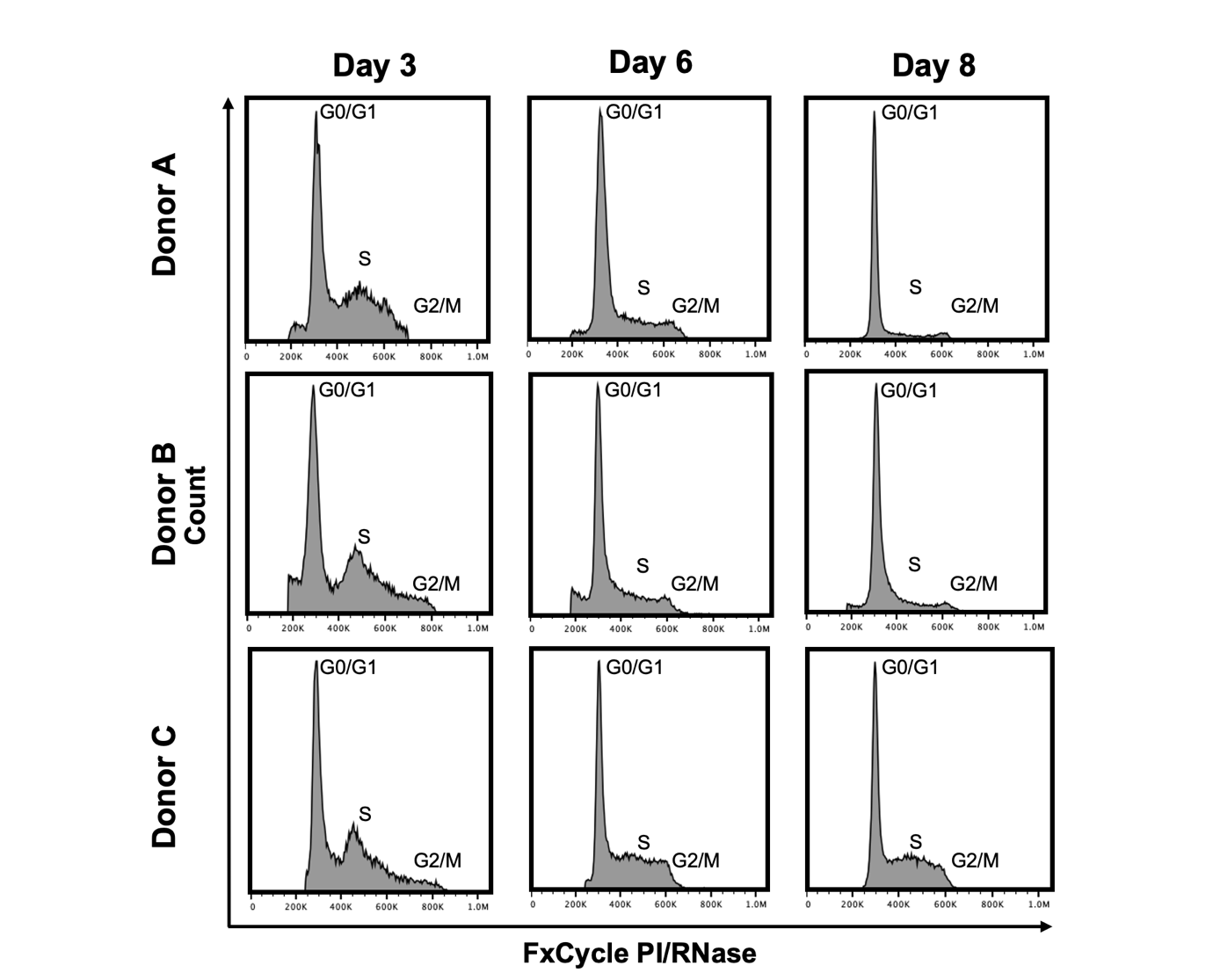

**Figure S8. Cell cycle analysis of human NK cells from three donors.** Cell cycle assay was performed on Days 3, 6, and 8 after activation with irradiated K562-mbIL15-41BBL feeder cells. DNA content stained with propidium iodide (PI) was analyzed by flow cytometry. The higher peaks on the left of histograms represent G0/G1 phase cells with 2n DNA content. The minor peaks on the right represent G2/M phase cells with 4n DNA content. Cells in S phase are in the middle with intermediate DNA content.

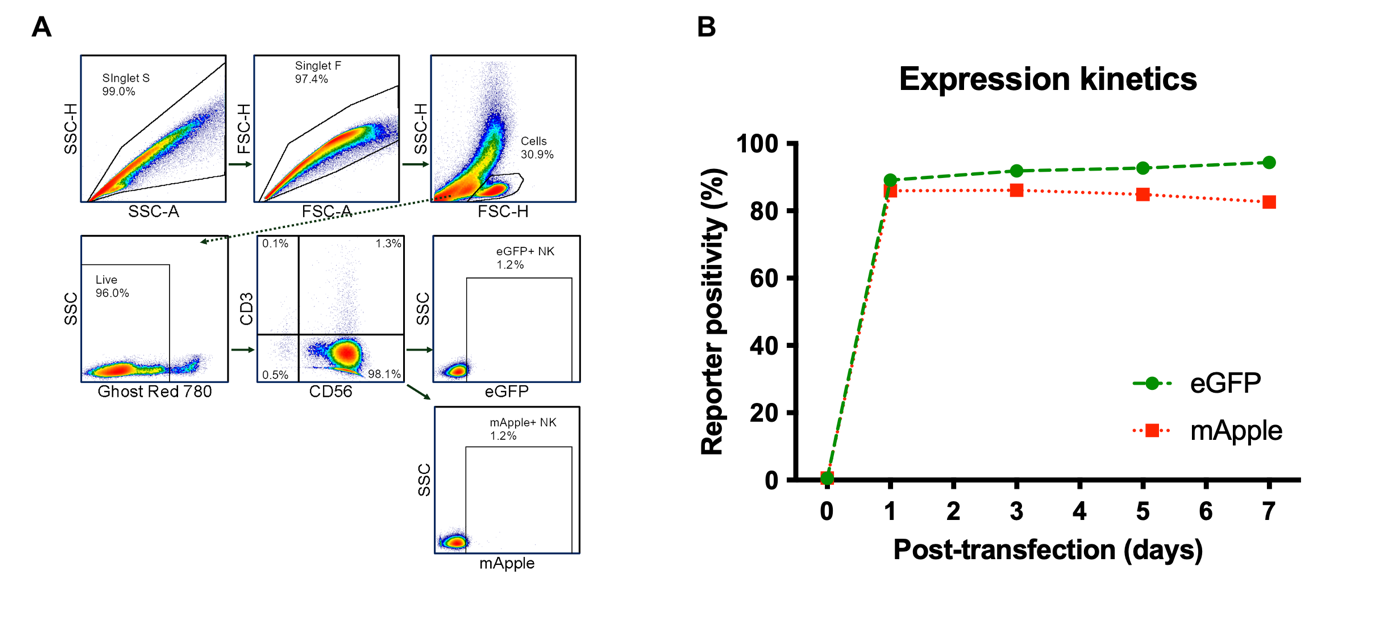

**Figure S9. Gating strategy and reporter expression kinetics.** (A) Representative gating strategy for viable CD3^-^CD56^+^ fluorescent reporter-positive NK cells. (B) The expanded NK cells were transfected with the GTA LNP (30 ng mRNA per 10,000 cells). eGFP and mApple expressed by the transfected NK cells from Donor A were monitored for one week using flow cytometry. Data are presented as mean (*n* = 3).

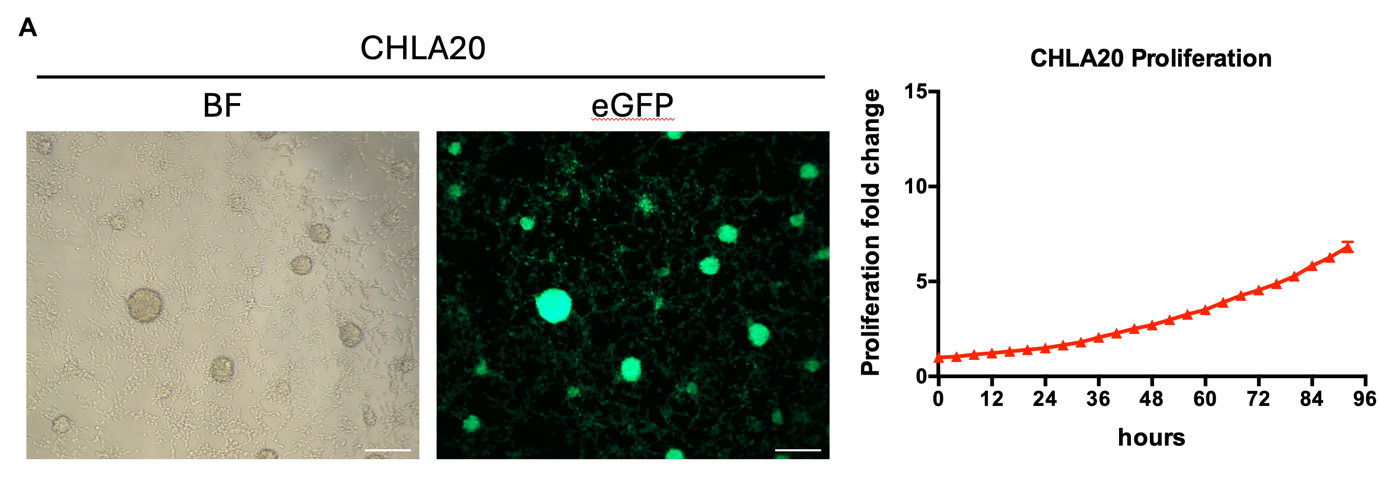

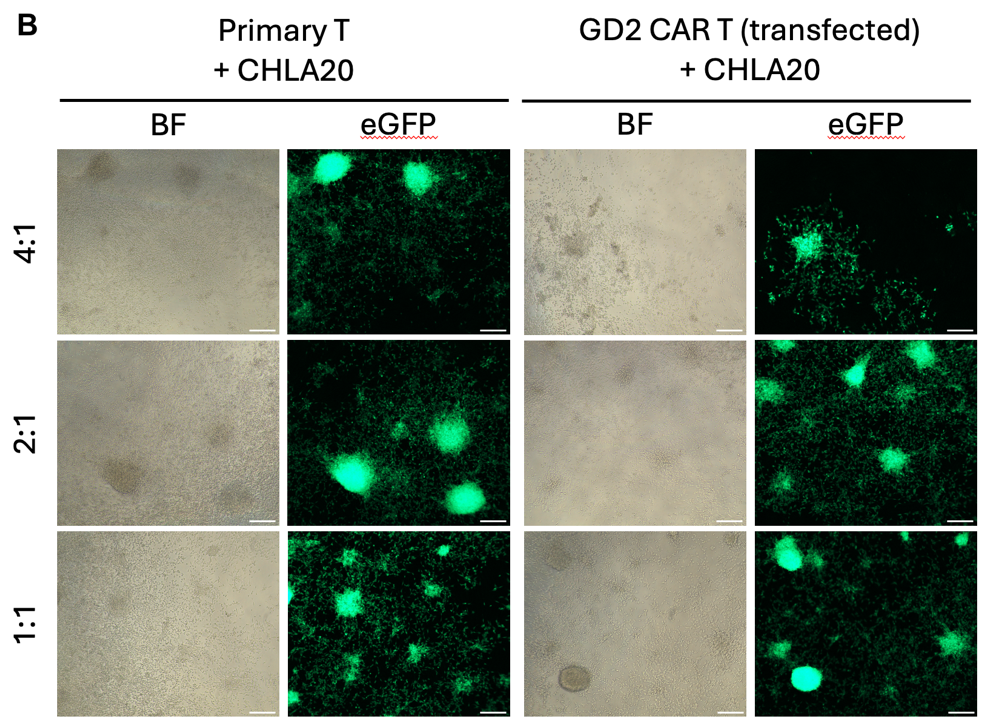

**Figure S10. IncuCyte assay using CHLA20 cells.** (A) Representative endpoint images of GFP^+^ CHLA20 cells. IncuCyte assay presented by proliferation fold change from 0 h to 92 h, recorded every 4 h. Data are presented as mean ± SEM (n = 3). Scale bar = 100 μm. (B) Representative endpoint images of GFP^+^ CHLA20 cells after co-culture with T cells or GD2 CAR T cells. Scale bar = 100 μm.
